# Fish environmental RNA sequencing sensitively captures accumulative stress responses through short-term aquarium sampling

**DOI:** 10.1101/2024.09.16.613370

**Authors:** Kaede Miyata, Yasuaki Inoue, Masayuki Yamane, Hiroshi Honda

**Affiliations:** R&D Safety Science Research, Kao Corporation, Ichikai–Machi, Haga–Gun, Tochigi 321-3497, Japan

**Keywords:** environmental RNA, RNA-sequence, biological monitoring, nature positive

## Abstract

The utility of environmental RNA (eRNA) in capturing biological responses to stresses has been discussed previously; however, the limited number of genes detected remains a significant hindrance to its widespread implementation. Here, we investigated the potential of eRNA to assess the health status of Japanese medaka fish exposed to linear alkylbenzene sulfonate. Analyzing eRNA and organismal RNA (oRNA) in aquarium water for > 12 h, we achieved high mapping rates and 10 times more differentially expressed genes than previously reported. This advancement has facilitated the previously unattainable capability of gene ontology (GO) analysis. In GO analysis, it was found that eRNA could detect nuclear genes in cellular component, and oRNA provided gene expression signatures in the short term, whereas eRNA demonstrated the potential to reflect cumulative gene expression signatures over a period of time in biological process. Moreover, eRNA exhibited high sensitivity in responding to genes associated with sphingolipid and ceramide biosynthesis, which are involved in inflammatory responses possibly originating from impaired cells. This finding aligns with the observations made in oRNA. Conclusively, eRNA-sequencing (eRNA-seq) using aquarium water emerges as a valuable high sensitivity tool for analyzing physiological stress. The findings of this study lay the foundation for further development of eRNA-seq technologies.

## 1. Introduction

As human populations expand,^1^ activities such as urbanization, industrial farming, resource overuse, and water pollution increase stress in freshwater ecosystems, possibly risking future biodiversity loss.^2,3^ Balancing environmental protection and economic growth is crucial.^4,5^ At the 15th Conference of the Parties to the Convention on Biological Diversity (COP15) in 2022, a plan termed “nature-positive” was adopted to reverse ecological decline, aiming to restore nature to the pre-2020 levels.^6^ To achieve these goals by 2030, there is an urgent need to develop new monitoring methods that are easier and more accurate than traditional field surveys in assessing biodiversity and stress conditions of inhabiting organisms.

Metabarcoding, using environmental DNA (eDNA) from aquatic environments, has revolutionized ecological surveys.^7–9^ This technique overcomes the limitations of traditional field surveys, such as high costs, lack of reproducibility, and invasiveness. Therefore, it is expected to play a key role in biodiversity assessments aimed at achieving nature-positive goals.^10, 11^ While eDNA metabarcoding provides information on species richness and abundance,^12–19^ it can lead to false positives as eDNA is stable and can persist in the environment for long durations.^20–22^ To address this issue, environmental RNA (eRNA), which degrades more rapidly than DNA,^23^ has been proposed as an alternative. Our previous studies have demonstrated the effectiveness of eRNA metabarcoding for ecological surveys with high predictive accuracy.^24–26^ Consequently, eRNA metabarcoding may facilitate simple and accurate ecological surveys, offering valuable insights into the presence or absence of habitants, which is crucial for biodiversity assessment.

However, detecting early-stage stresses involved in population decline or extinction through eDNA/RNA metabarcoding remains challenging. Early identification of stresses that adversely affect habitat organisms can benefit effective ecosystem conservation efforts.

Recently, research on “environmental transcriptomics,” which captures the stress state using transcripts (messenger RNA; mRNA) in eRNA using RNA-sequencing (RNA-seq) has commenced .^27^ Such studies performed in aquaria have suggested that environmental micro RNAs (miRNA) and mRNAs can act as indicators for understanding the stress state of organisms in water.^28–30^ However, the number of differentially expressed environmental miRNAs or mRNAs in water tends to be very low. Therefore, understanding gene expression profiles of eRNA in response to stimulation remains limited due to the absence of a detailed gene ontology (GO) enrichment analysis. One of the reasons for the low number of genes detected in water is the difficulty in controlling the activity of microorganisms in the aquarium. Typically, the number of microorganisms in aquaria tends to increase during prolonged experiments, leading to the degradation of eDNA and eRNA.^31^ As mapping degraded eRNA to the reference sequence is challenging, the limited detection of differentially expressed genes (DEGs) from water samples may be attributed to eRNA degradation. Therefore, sequencing mRNA sequences in water before degradation is crucial for accurately capturing stress information from eRNA released by aquatic organisms. To enhance the mapping rate, we focused on optimizing the sampling time. By conducting environmental RNA sequencing (eRNA-seq) analyses with a high mapping rate, we aimed to capture the stress state of organisms in aquarium water by evaluating the relationship between the mapping rate and sampling time. It may be possible to validate whether eRNA can effectively capture changes in gene expression in aquatic organisms and assess the usefulness and limitations of the technology for real-environmental applications through environmental transcriptome analysis with a high mapping rate.

In this study, we conducted RNA-seq analyses of organismal RNA (oRNA) from Japanese medaka fish (*Oryzias latipes*; *O. latipes*) exposed to linear alkylbenzene sulfonate (LAS), along with eRNA from their aquarium water at suitable sampling times during LAS exposure. LAS, a surfactant commonly present in daily-consumer products and industrial detergents, induces oxidative stress in exposed fish by generating reactive oxygen species (ROS).^32,33^ The aims of this study are to (1) develop eRNA-seq with enhanced analytical capabilities in aquarium settings and (2) verify whether the stress response of individuals due to LAS exposure can be detected using the aquarium water. This study aims to demonstrate that eRNA monitoring can provide insights not only into the presence or absence of organisms but also into their stress state, offering valuable information for effective biodiversity conservation planning.

## 2. Materials and methods

### 2.1. Experimental design

The experimental design is illustrated in Fig. S1. The appropriate sampling time was predetermined through careful observations of the dynamics of eRNA release from *O. latipes* and its degradation by bacteria over time to assess their impact on the mapping rate. The appropriate LAS concentration was determined by monitoring the relationship between the state of *O. latipes* and LAS concentration. These experiments were conducted with *O. latipes* in the aquarium.

Detailed methodologies for these trials, aimed at identifying the appropriate sampling time and LAS concentration are elaborated in Texts S1, S2, and S3. The sampling times and LAS concentrations determined through these experiments were used to conduct the main experiment. The methods used for the main experiment are described in detail in the following sections.

Prior to the experiments, the glass aquaria (length, width, and height: 20 × 15 × 20 cm) were cleaned using detergent, 10% chlorine bleach solution, and ethanol, followed by multiple rinses with distilled water to remove any nucleic acid.

The exposure concentrations included control and LAS-treated groups at 9 mg/L. Although these exposure concentrations are considerably higher than those typically found in rivers,^34^ the 9 mg/L level was chosen because it is the lowest concentration that affects by *O. latipes* (Table S1). This setting was used to observe the effects on *O. latipes* in the dose range finding study.

The glass aquaria were filled with 3 L of LAS solution and were arranged on the water bath. The water in the glass aquaria was maintained at 24 ± 1 °C using an open circuit chiller (Taitec Corp., Saitama, Japan) under a 16-h light: 8-h dark cycle and continuously aerated using glass tubes and air pump. Each concentration had three replicates, and each replicate contained fifteen fish. The *O. latipes* strain, NIES_R, utilized in our experiment was obtained from the National Institute for Environmental Studies (Japan) and has been reared in our laboratory for over two generations. Fish were maintained in consistent conditions (24 ± 1 °C and a photoperiod of 16 h-light: 8 h-dark). They were fed brine shrimp twice daily. We used 3–4-month-old adult male fish with a similar body size (2.28 ± 0.118 cm and 0.16 ± 0.024 g; Table S2). Similarly, a negative control (a glass aquaria with no fish but dechlorinated water) was prepared to confirm no contamination of RNA. No food was given to the sample fish from 24 h before the start of the experiment until its end to control the effect of feces. The water temperature, pH, and DO were measured at each sampling time point using a digital thermometer SN3000 (NETSUKEN Corp., Tokyo, Japan) and pH/DO meter D-55 (HORIBA, Kyoto, Japan). The employed LAS analysis methods are detailed in Text S4.

At each time point, water samples and fish were collected at 0, 3, 6 and 12 h. All fish were removed and anesthetized immediately on ice. After removal of 15 fish, 2 L water samples were filtered through a Sterivex cartridge (nominal pore size, 0.45 µm; Millipore, Billerica, MA, USA) using a Masterflex peristaltic pump (Cole-Parmer, Vernon Hills, IL, USA). RNA extraction from the Sterivex cartridge was performed immediately.

All animal experiments were performed according to the guidelines of the Experimental Animal Facility of Kao Corporation’s R&D Department.

### 2.2. Sample preparation

The total RNA extraction of water samples and fish were performed using the RNeasy Plus Universal Mini Kit (Qiagen, Hilden, Germany) and treated with DNase using RNase-Free DNase Set (Qiagen) to remove contaminating genomic DNA. RNA was extracted from whole fish bodies that were randomly selected fish (n = 5) and collected at each sampling point according to the manufacturer’s protocol. To reduce the variation among individuals, we combined equal amounts of RNA from five fish in the same aquarium to create one sample. This sample was then divided into three subsamples per aquarium (3 aquaria × 3 subsamples × 2 LAS concentration (with or without LAS)). eRNA was extracted from the water sample with some modifications of the manufacturer’s protocol. Briefly, the QIAzol lysis reagent (Qiagen) was introduced into the Sterivex cartridge to lyse the tissue contained in the filter. The cartridge was then incubated using mild rotation at 25 °C for 60 min. The QIAzol lysis reagent was collected by centrifuging. Then 100 µL gDNA Eliminator Solution was added to the lysis reagent and vortexed for 15 s.

The subsequent steps were conducted according to the manufacturer’s protocol. RNA concentration and quality were measured using Qubit 4.0 Fluorometer (Life Technologies, CA, USA) and Agilent 2200 TapeStation (Agilent Technologies, Inc., CA, USA), respectively. cDNA synthesis was performed with SMART-Seq mRNA (TaKaRa Bio Inc., Shiga, Japan) using 30 ng RNA extracted from whole fish or water samples as input according to the manufacturer’s protocol. cDNA was purified using AMPure XP beads (Beckman Coulter, Brea, CA, USA). Next, 100 ng cDNA from each sample was used to generate a sequence library using Illumina DNA Prep kit with IDT^®^ for Illumina^®^ DNA/RNA UD Indexes (Illumina, San Diego, CA, USA) following the manufacturer’s protocol. We prepared a total of 29 sequencing libraries, which consisted of three replicated samples, four sampling time points, two treatment groups, and five quality controls. Subsequently, the two pools of libraries obtained from both whole fish and water samples were sequenced individually on a Next Seq 2000 (Illumina) using the P2 flow cell (Illumina) as the sequencing reagent. This sequencing process resulted in obtaining paired-end reads with a length of 100 base pairs (2 × 50 bp).

### 2.3. Data analysis

The sequenced raw reads were trimmed using Trimmomatic ver. 0.39^35^ with the following parameters: ILLUMINACLIP:adapters.fa:2:30:10, LEADING:3, TRAILING:3, SLIDINGWINDOW:4:15, MINLEN:36 to remove low-quality reads and adaptors. The quality control of reads were performed using FastQC ver 0.11.9 with default setting.^36^ We aligned clean reads to the reference genome of *O. latipes* (accession number: GCF_002234675.1, accessed on February 28, 2023) using BWA ver. 0.7.17 with default setting,^37,38^ and subsequently calculated mapping rates. Mapping rates were calculated as the number of reads mapped to the reference database per total paired reads in sequencing. The read counts were quantified using SAMtools ver. 1.10 with default setting.^39^ Genes with read counts of < 10 were excluded from the Microsoft Excel dataset. The read counts were normalized using trimmed mean of M values (TMM) method^40,41^ into edgeR ver. 3.15 package^41,42^ by means of the calcNormFactors function. DEGs were determined using R software ver. 4.2.2 and edgeR ver. 3.15 based on the criteria of a false discovery rate (FDR) < 0.05 and fold change ≥ 1.1 (for oRNA) or 2 (for eRNA).^41,42^ DEGs were subjected to gene enrichment analysis using DAVID ver. 2021^43^ with the complete *O. latipes* and *Danio rerio* (zebrafish) genomes as the background. DAVID is an online analytical tool employed for the scientific extraction of biological implications from genes or proteins.

DAVID was used for CC and BP annotations, employing settings with thresholds set at Count 2 and EASE score 0.1. The Fisher’s exact test in DAVID was used to calculate *p* values; *p* values < 0.05 were considered statistically significant.

### 2.4. Quantification of genes expressions related to inflammation in oRNA and eRNA using ddPCR

ORNA and eRNA samples were quantified using a Bio-Rad QX200 AutoDG Droplet Digital PCR system (Bio-Rad Laboratories, Hercules, CA, USA), including negative filtration, extraction, and DNA synthesis controls, to precisely examine quantitative differences in oRNA and eRNA. Before performing ddPCR analysis, cDNA samples were diluted to levels between 1:100 and 1:10 to reduce inhibition and ensure that they were within the measurement range of ddPCR. PCR was performed in a 22 μL reaction volume containing 11 μL of 2 × ddPCR EvaGreen Supermix, 1.1 μL of template cDNA, 0.3 μL of primer pairs (10 μM), and 9.3 μL of sterile deionized water. Thermal cycling was performed at 95 °C for 5 min; 40 cycles of 95 °C for 30 s and by setting the temperature of each primer pairs for 1 min; and 4 °C for 5 min; 95 °C for 5 min (Tables S3 and S4). Each sample was quantified in triplicate, with negative (sterile distilled water) and positive controls (the total cDNA extracted from *O. latipes*; 0.2 ng/μL) included in each plate. Data were analyzed using the Bio-Rad QuantaSoft software version 1.7.4. Samples with < 10,000 accepted droplets were remeasured until 10,000 accepted droplets were achieved. The threshold of cDNA for droplet positivity was automatically set.

### 2.5. Statistical analyses

The F-test was used to determine the homogeneity of variance across log-scale data of read counts and copy numbers. For comparing read counts and copy numbers, a two-tailed paired *t*- test was performed when the underlying distributions were parametric and the variance was homogeneous, whereas the Welch *t*-test was performed when the variance was heterogeneous. The −log *p* value of the GO analysis on BP was used to evaluate the difference of the enriched functional terms. PCA was performed based on the number of read counts using OriginPro 2021b (OriginLab, Northampton, MA, USA). All statistical analyses, except the Welch *t*-test, hierarchical clustering, and the validation of read counts/copy numbers, were performed using OriginPro 2021b. The Welch *t*-test was performed using R (ver. 4.3.1). Hierarchical clustering analysis and the validation of read counts/copy numbers were performed using ComplexHeatmap and fdrtool, packages, respectively, in R (ver. 4.3.1). Venn plots with four circles, balloon plots, and heatmap with hierarchical clustering were illustrated using R (ver. 4.3.1). Circle plots, bar plots, heatmap, and PCA plot were illustrated using OriginPro 2021b. Statistical significance for differential expression, excluding multiple testing, was determined at a threshold of *p* < 0.05. Multiple testing correction employed FDR-adjusted *p* (*q*) < 0.1.

## 3. Results

### 3.1. Finding appropriate sampling time and LAS concentration

During the determination of the optimal sampling time, it was found that the initial mapping rate of eRNA was 11.4–20.1%, decreasing to < 1% after ≥ 24 h (Table S5). This decline in both eRNA concentration and mapping rate suggests potential eRNA degradation facilitated by microbial activity within the closed system of the aquarium, in contrast to the open system of the real environment.

In an investigation on determining the optimal sampling time for assessing *O. latipes* eRNA and bacterial DNA levels in aquarium water, it was observed that following the introduction of *O. latipes* into the aquarium, the eRNA concentration steadily increased until reaching a peak at 6 h. However, beyond this point, eRNA levels declined, with concentrations at 24 h notably lower than those at the initial time point (10 min), despite the continued presence of live *O. latipes* (Fig. 1A and Table S3). The bacterial growth curve data aligned well with the logistic population growth model, indicating a gradual increase in bacterial DNA over time, with a projected plateau at approximately the 24-h mark (Fig. 1B and Table S3). Consequently, water sampling was limited to within the first 12 h to minimize the impact of RNA degradation on eRNA-seq analyses.

**Fig. 1.**
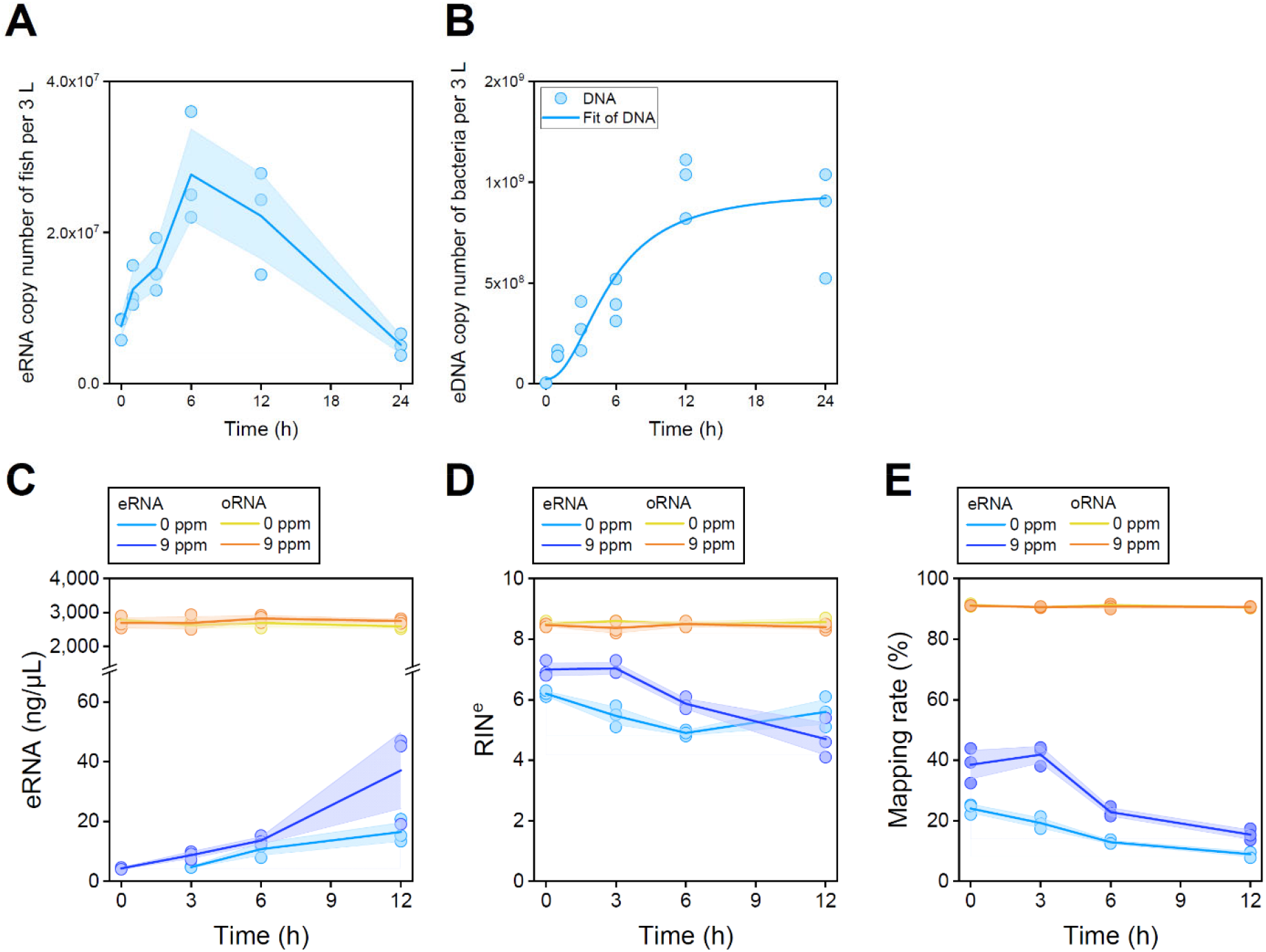
Time-dependent changes in properties of eRNA and bacterial abundance in the aquarium. (**A**) Time-dependent change in the eRNA copy number of *O. latipes*. The plots represent three replicates and the indicates their mean value. The light blue shaded area represents the standard deviation. (**B**) Time-dependent changes in the eDNA copy number of bacteria. The points show the three replicates, and the line represents the logistic model fit, with *R*^2^ = 0.82. Time- dependent changes in eRNA (**C**) concentration, (**D**) quality, and (**E**) mapping rate. RNA quality was represented as the Integrity Number equivalent [RIN^e^]. The points and lines show the three replicates and their mean value, respectively, with the areas shaded in color indicating standard deviations.

During the determination of the optimal LAS concentration, it was found that all *O. latipes* were alive in the control and LAS 8 mg/L groups over 0–24 h, and 4 out of 15 and 10 out of 15 *O. latipes* were dead at LAS 9 mg/L and 10 mg/L groups at 24 h, respectively (Table S1).

Consequently, LAS 9 mg/L was used in the main experiment, which was the lowest concentration that affected *O. latipes*.

### 3.2. Summary of the main aquarium experiment for RNA-seq

The concentrations of LAS during 0-12 h are summarized in Table S6 (in Text S4). The effects of LAS exposure to *O. latipes* and water quality are summarized in Table S7. One out of 15 *O. latipes* subjected to 9 mg/L of LAS exposure for 12 h was dead in two of the three glass aquaria. Temperature, dissolved oxygen (DO), and pH did not change significantly during the experiment.

The concentration and quality (that is, RNA Integrity Number equivalent [RIN^e^]) were measured using Qubit Fluorometer and TapeStation, respectively. The concentration and quality of oRNA were almost the same with or without LAS exposure (Figs. 1C, D, and E and Table S8). The concentrations of oRNA were higher than those of eRNA and stable throughout the experiment. Raw reads and mapping rates (90.5–91.2%) were similar among the groups of oRNA. In contrast, the concentration of eRNA exposed to LAS increased with time, while RIN^e^ decreased; the concentration of eRNA without LAS increased slightly and RIN^e^ decreased up to 6 h and increased slightly at 12 h. Raw reads were similar among the groups of eRNA, but mapping rates decreased with time (9.0–41.9%). The mapping rate of eRNA increased with LAS exposure.

### 3.3. Gene detection of *Oryzias latipes* and analysis of sources

Sequencing eRNA extracted from aquarium water and oRNA obtained from *O. latipes* tissues at 0 h without LAS identified 9,854 (38.2%) and 22,173 (86.0%) genes, respectively, out of the 25,795 genes cataloged for *O. latipes* in the database (Fig. 2A). Notably, 9,811 (99.6%) of the genes detected in eRNA were also identified in oRNA. The oRNA read counts correlated with those of eRNA (*r* = 0.56, *p* < 0.01; Figs. 2B and S2) and most genes were distributed along the 1:1 line. However, some genes were detected using either oRNA or eRNA analysis. To analyze the distribution of gene expression, the GO analysis on the cellular component (CC) was conducted (*p* < 0.05; Figs. 2C and D). Of the 12,362 genes detected only in oRNA, 4,253 genes were annotated. Among the annotated genes, 3,543 (83.3%), 615 (14.5%), and 95 (2.2%) genes were derived from extracellular, intracellular, and both sources, respectively (Fig. 2C, left).

**Fig. 2.**
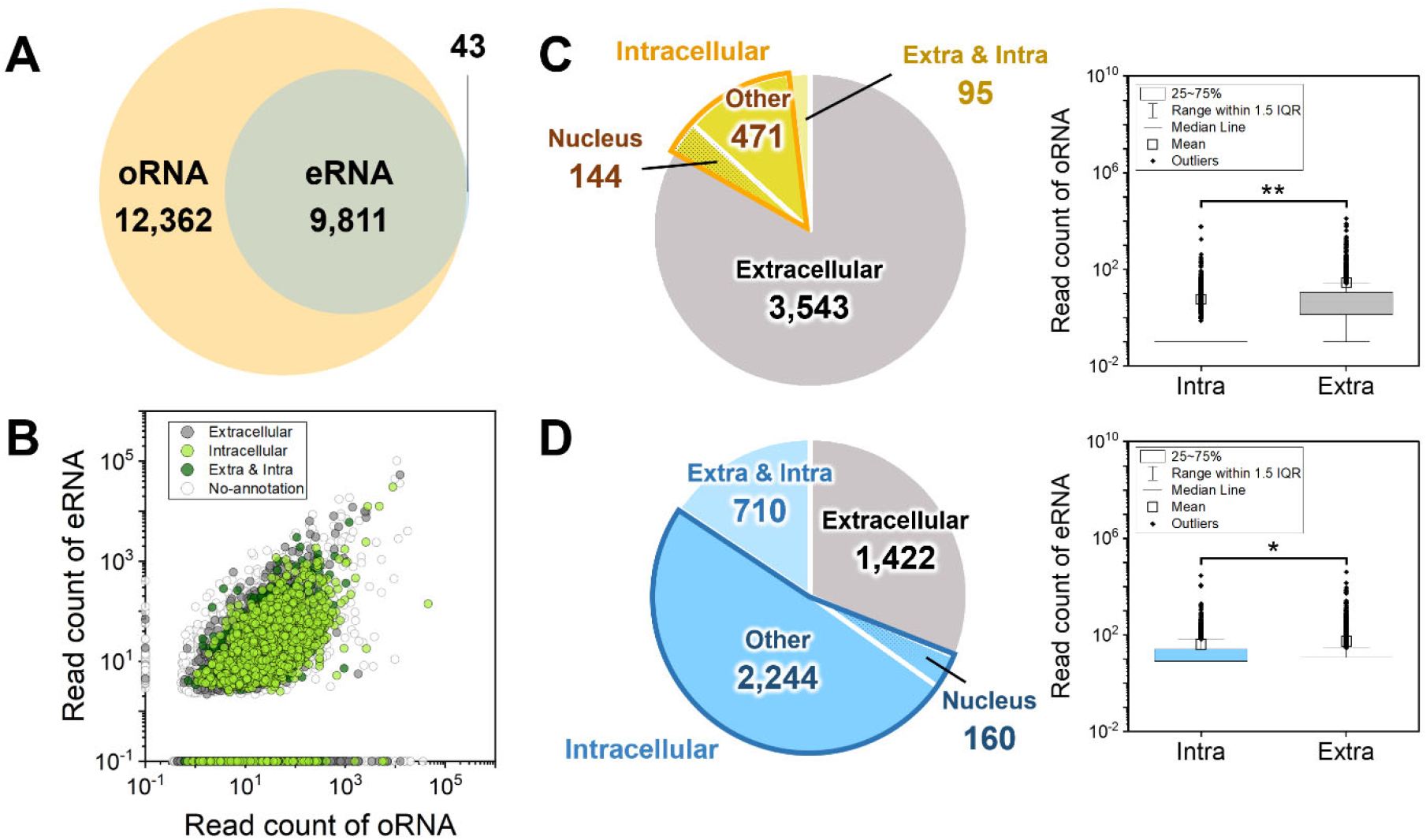
Comparison of expressed genes between oRNA and eRNA at 0 h without LAS in the GO analysis on cellular component (CC), respectively (*p* < 0.05). (**A**) Venn diagram showing the number of genes detected in oRNA and eRNA. (**B**) Scatter plot showing read counts of all genes between oRNA and eRNA on the GO analysis on CC (*p* < 0.05). Genes are categorized as extracellular (gray dots), intracellular (light green dots), both (green dots), and no-annotation (white dots). Pie charts showing the number of genes detected only in (**C**) oRNA or (**D**) both oRNA and eRNA. The gray, yellow (or blue), and light yellow (or light blue) areas indicate genes derived from extracellular, intracellular, and both sources, respectively. Box plots for comparison of intracellular or extracellular sources. Orange or blue box: intracellular; Gray box: extracellular. \**p* < 0.05. \*\**p* < 0.01.

Localization to intracellular organelles, such as the nucleus, was categorized as intracellular, while all other localizations were considered extracellular. Among the 615 genes derived from the intracellular source, 144 (23.4%) genes were derived from the nucleus. Of the 9,811 genes detected in both oRNA and eRNA, 4,536 genes were annotated. Among the annotated genes, 1,422 (31.3%), 2,244 (49.5%), and 710 (15.7%) genes were derived from extracellular, intracellular, and both sources, respectively (Fig. 2D, left). Among the 2,404 genes derived from the intracellular source, 160 (6.7%) genes were derived from the nucleus. No gene among the 43 genes detected only in the eRNA was annotated.

The distribution of oRNA read counts derived from the extracellular source was significantly higher than that derived from the intracellular source (Welch *t*-test; *p* < 0.01; Fig. 2C, right). However, the distribution of eRNA read counts derived from the intracellular source was significantly higher than that derived from the extracellular source (Welch *t*-test; *p* < 0.01; Fig. 2D, right). Therefore, our method for obtaining eRNA including intracellular genes enable the capture of intracellular gene expression signature including the nucleus.

Genes detected in both oRNA and eRNA at 0 h without LAS were subjected to GO for biological process (BP) identification. In oRNA (12,362 genes), 83 functional terms were identified, while oRNA/eRNA (9,811 genes) revealed 76 functional terms, with no overlap between the two sets (Fig. S3). The top five enriched terms in oRNA (12,362 genes) included visual perception (GO:0007601), rRNA processing (GO:0006364), negative regulation of endopeptidase activity (GO:0010951), meiotic cell cycle (GO:0051321), and tRNA wobble uridine modification (GO:0002098). The top five enriched terms in oRNA/eRNA (9,811 genes) included ubiquitin-dependent protein catabolic process (GO:0006511), intracellular protein transport (GO:0006886), ER to Golgi vesicle-mediated transport (GO:0006888), protein polyubiquitination (GO:0000209), and mitophagy (GO:0000422) (Table S9).

### 3.4. Differential gene expression and functional annotation effected by LAS in oRNA and eRNA

To assess the impact of LAS exposure on *O. latipes*, we compared the expressed genes in oRNA and eRNA with or without LAS (Fig. S4). At 0 (10 min), 3, 6 and 12 h after exposure to LAS, oRNA exhibited the upregulation of 549, 180, 1,102, and 1,783 DEGs, and downregulation of 678, 187, 1,125, and 1,308 DEGs (Fig. S4B). In parallel, eRNA showed the upregulation of 89, 710, 1,082, and 1,189 DEGs, and downregulation of 458, 1,082, 889, and 1,816 DEGs, respectively (Fig. S4D).

The comparative analysis between samples without and with LAS for the whole transcriptome revealed characteristic changes in functional terms derived from DEGs during the 0–12 h period in oRNA (Fig. 3A and Table S10) and eRNA (Fig. 3B and Table S10). In oRNA, almost all enriched functional terms were detected at each time point, with a notable trend observed for upregulated DEGs, indicating a point-by-point enrichment pattern. In eRNA, functional terms enriched at any time point were frequently detected in later time points (e.g., functions detected at 0 h (10 min) were frequently detected at 3, 6, and 12 h), indicating that eRNA exhibits a superior capability to capture cumulative gene expression signatures throughout the experiment. Although the cumulative effect was less apparent in the downregulated genes (Fig. 3B, right), the degradation of genes by microorganisms in water may diminish the visibility of their effects.

**Fig. 3.**
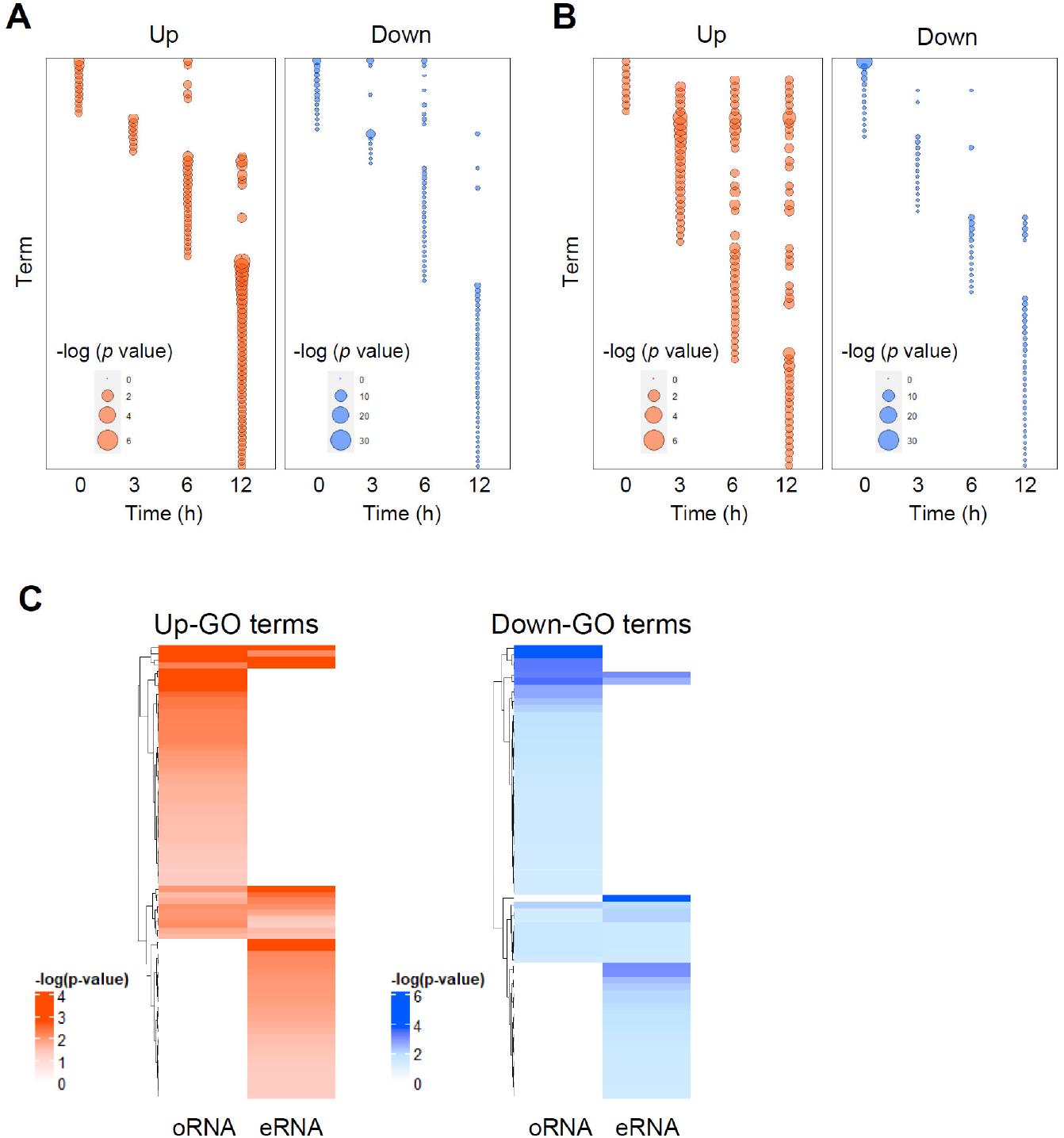
Functional terms in oRNA and eRNA between LAS treatment groups and control groups. Balloon plots illustrate the relationship of −log *p* values for functional terms in the GO analysis on BP in (**A**) oRNA or (**B**) eRNA (*p* < 0.05). The vertical axis lists the upregulated (left) and downregulated (right) functional terms, which are merged from each time point. Larger dots represent higher values. (**C**) Heatmap showing the relationship of −log *p* value in functional terms annotated from (right) merged-upregulated-DEGs and (left) merged-downregulated-DEGs in the GO analysis on BP in oRNA and eRNA.

Heatmap clustering analysis was performed to reveal the differences in GO terms related to BPs that are enriched in oRNA and eRNA. Although there was no overlap in functional terms enriched by oRNA and eRNA in the original expression (at 0 h (10 min) without LAS; Fig. S3), the heatmap of GO terms enriched by up- and downregulated genes in LAS-treated groups on the BP annotation chart in oRNA and eRNA clearly show the separation between oRNA, eRNA, and both (Fig. 3C and Table S11). Among the 77 functional terms enriched in merged-up-DEGs, combining up-DEGs at 0, 3, 6 and 12 h, in oRNA and eRNA, 13 (16.9%) up-terms were identified in both oRNA and eRNA (Fig. 3C left and Table S11). Among the 13 up-terms, signaling pathways such as sphingolipid biosynthetic process (GO:0030148; e.g., *elovl1a*, *elovl7a*, *degs1*, *degs2*, *LOC101172187*, and *fa2h*), epidermal growth factor receptor (EGFR) signaling pathway (GO:0007173; e.g., *epgn*, *LOC101166137*, *hbegfa*, and *src*), and ceramide biosynthetic process (GO:0046513; e.g., *degs1* and *degs2*) related to the inflammatory response, autophagy, and apoptosis were included. Among 67 functional terms enriched on merged-down- DEGs, which were merged with each down-DEG at 0, 3, 6 and 12 h, in oRNA and eRNA, 11 (16.4%) down-terms were identified on both oRNA and eRNA (Fig. 3C right and Table S11).

Among the 11 down-terms, signaling pathways such as defense response (GO:0006952; e.g., *LOC101174108*, *LOC101168657*, and *LOC101174598*), antigen processing and presentation (GO:0019882; e.g., *cd74a*, *LOC111946286*, and *LOC101163710*), and immune response (GO:0006955; e.g., *ctsl.1*, *cd74a*, and *zgc:174863*) were included.

The characteristics of GO terms detectable in oRNA, eRNA, and both were further analyzed, respectively. In GO terms in the BP annotation chart in oRNA (*p* < 0.05), the signaling pathways in response to oxidative stress were activated by the exposure to LAS (Table S11).

Genes involved in the xenobiotic metabolic process (GO:0006805; e.g., *LOC101156566*, *LOC101159216*, *LOC101157767*, *cyp2p3*, *LOC111948423*, *LOC101156965*, *LOC101164878*, *LOC105353996*, *ahrr*, and *LOC101157528*), in positive regulation of I-kappaB kinase/NF-kappaB (NF-κB) signaling (GO:0043123; e.g., *trim13, LOC105356874, tnfrsf19, ror1, traf3ip2l, tab2, rbck1, LOC105355953,* and *faslg*), and in negative regulation of mitogen-activated protein kinase (MAPK) cascade (GO:0043409; e.g., *dusp 1, 5*, *6*, *8a*, *10* and *13, pebp1, LOC101175354,* and *LOC101173884*) were downregulated, and in the glutathione metabolic process (GO:0006749; e.g., *ethe1*, *clic1*, *clic3*, *LOC101171103*, *gpx1b*, *gsr*, *LOC101161407*, *mars1*, *LOC105354455*, and *LOC101175403*), in sphingolipid biosynthetic process (GO:0030148; e.g., *hacd2*, *elovl1a*, *elovl5*, *elovl7a*, *degs1*, *degs2*, *LOC101172187*, and *fa2h*), in ceramide biosynthetic process (GO:0046513; e.g., *cers1*, *cers2b*, *cers3b*, *LOC101166550*, *degs1*, *degs2*, *sgms1*, and *LOC101167981*), and in autophagy (GO:0006914; e.g., *ulk1a, zgc:92606, fbxo32, ulk2, gabarapl2, usp10, gabarapa, wdr45, atg9b, vmp1, atg101, si:ch211-153b23.5, map1lc3b, atg4b,* and *vcp*) were upregulated (Fig. 4 and Table S12). Other functions, such as cytokine-mediated signaling pathway (GO:0019221; e.g., *lepr*, *stat6*, *il13ra2*, *f3a*, *LOC105357584*, *ifngr1*, *LOC101164128*, *stat3*, *ghra*, *LOC101169209* (*IL-1β*), and *prlrb*) were upregulated. The gene expression of *IL-1β*, one of the most representative genes of the inflammatory response and immune system, using droplet-digital PCR (ddPCR) were also upregulated (Fig. S5 and Table S4).

**Fig. 4.**
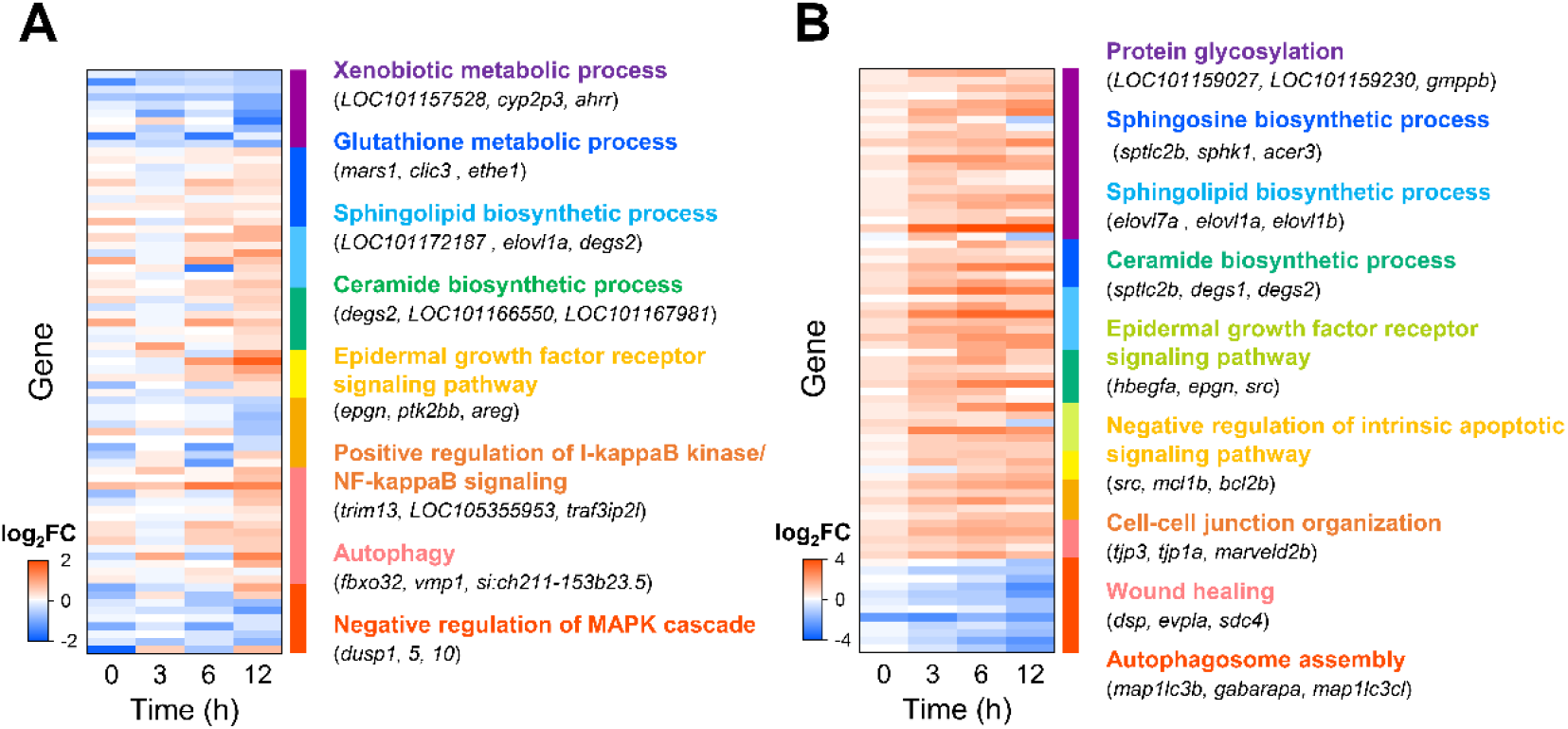
Functional terms related to inflammatory response in oRNA and eRNA between LAS treatment groups vs control groups. Heatmap of DEGs in each functional terms in the GO analysis on BP at each time point in (**A**) oRNA and (**B**) eRNA. Color represents −log *p* value of each gene. Red or blue indicate up- or downregulated, respectively. White indicates no differential expression. log2FC; log2 fold change.

In the BP annotation chart in eRNA (*p* < 0.05), GO terms related to the inflammatory response and immune system, such as protein glycosylation (GO:0006486), sphingosine biosynthetic process (GO:0046512), sphingolipid biosynthetic process (GO:0030148), ceramide biosynthetic process (GO:0046513), and wound healing (GO:0042060), were significantly upregulated although the series of the signaling pathways activated during oxidative stress was not detected in eRNA. In addition, the functional term, wound healing, was significantly enriched in eRNA. As these genes are potential stress markers in water, their expression levels in oRNA and eRNA were measured using ddPCR. The ddPCR results of these wound healing-related genes, such as *dsp*, *evpla*, *evplb*, and *ppl*, were consistent with those of the RNA-seq analysis (Fig. S6 and Table S4). Contrary to expectations, some GO terms associated with inflammation and immunity, such as autophagosome assembly (GO:0000045), tend to be downregulated, despite the expectation of their upregulation in oxidative stress-related pathways.

### 3.5. Gene expression patterns revealed by principal component analysis (PCA) and important discriminatory genes

PCA was employed to examine genes upregulated in eRNA, aiming to identify commonly responsive genes to LAS exposure in both oRNA and eRNA. The analysis focused on 1,685 DEGs detected in eRNA and oRNA. The resultant PCA plot (Fig. 5) shows distinct separation among experimental groups. The primary separation along the first principal component (PC1), accounting for 53.1% of the overall variance, reflected the RNA type. The second principal component (PC2), representing 20.2% of the overall variance and associated with LAS exposure, revealed distinct clustering of samples. Genes with significant loading scores for PC2 included *LOC101175403, mars1, LOC105354455, and clic1* in the glutathione metabolic process (GO:0006749)*, elovl1a, fa2h* and *elovl7a* in the sphingolipid biosynthetic process (GO:0030148)*, degs1* in the sphingolipid biosynthetic process (GO:0030148) and ceramide biosynthetic process (GO:0046513)*, epgn, LOC101166137, hbegfa,* and *src* in the EGFR signaling pathway (GO:0007173)*, vcp* in autophagy (GO:0006914), *dusp1* and *dusp5* in the negative regulation of MAPK cascade (GO:0043409), and *dsp*, *evpla* and *evplb* in the wound healing (GO:0042060). These genes were consistently identified as DEGs between oRNA and eRNA; they were corroborated by functional analysis using GO, indicating their potential as biomarkers for characterizing LAS exposure. Collectively PC1 and PC2 explained 73.3% of the total variance of all principal components. The response to LAS at 12 h appeared to be similar for oRNA and eRNA, although time dependence was not consistently evident.

**Fig. 5.**
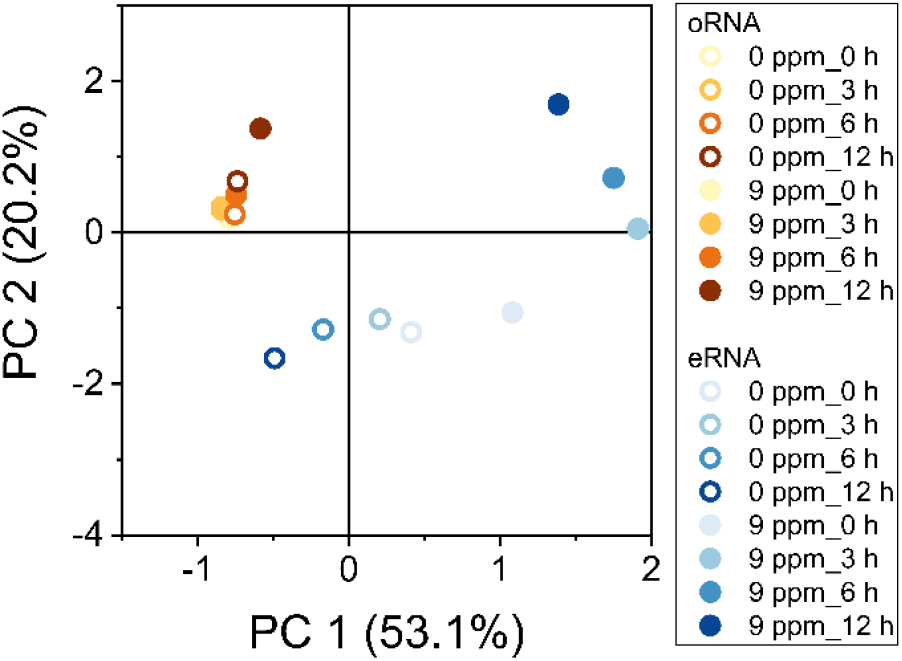
Distribution of gene expression patterns revealed by PCA. Principal component analysis (PCA) based on read counts of 1,685 upregulated DEGs in eRNA between LAS treatment groups vs control groups. PC 1 and PC 2 explained 53.1 and 20.2% of data variance, respectively. Different colors and symbols represent with or without LAS treatments, RNA sources and time points, respectively.

### 3.6. Validation of eRNA-seq data using reverse transcription-droplet digital PCR

Based on the functional analysis and PCA, genes associated with the sphingolipid and ceramide biosynthetic process emerged as potential stress markers. To verify the RNA-seq findings, we employed reverse transcription-droplet digital PCR (RT-ddPCR) on un-sequenced samples (Table S4). Fourteen genes from oRNA and thirteen genes from eRNA, identified as DEGs and linked to sphingolipid and ceramide biosynthetic processes, were subjected to validation.

Notably, genes such as *elovl1a*, *degs1*, and *fa2h*, which were identified as DEGs associated with LAS effects (as indicated by PC2 in the PCA analysis), were validated through ddPCR. The ddPCR results were consistent with those of the RNA-seq analysis (Fig. 6). Copy numbers of *elovl1a*, *degs1*, and *fa2h*, significantly altered by LAS, mirrored the read counts at the next- generation sequencing (NGS). Overall, the changes relative to the control group tended to be more pronounced for eRNA than for oRNA, suggesting that oRNA and eRNA potentially have distinct subsets of genes with varying detectability.

**Fig. 6.**
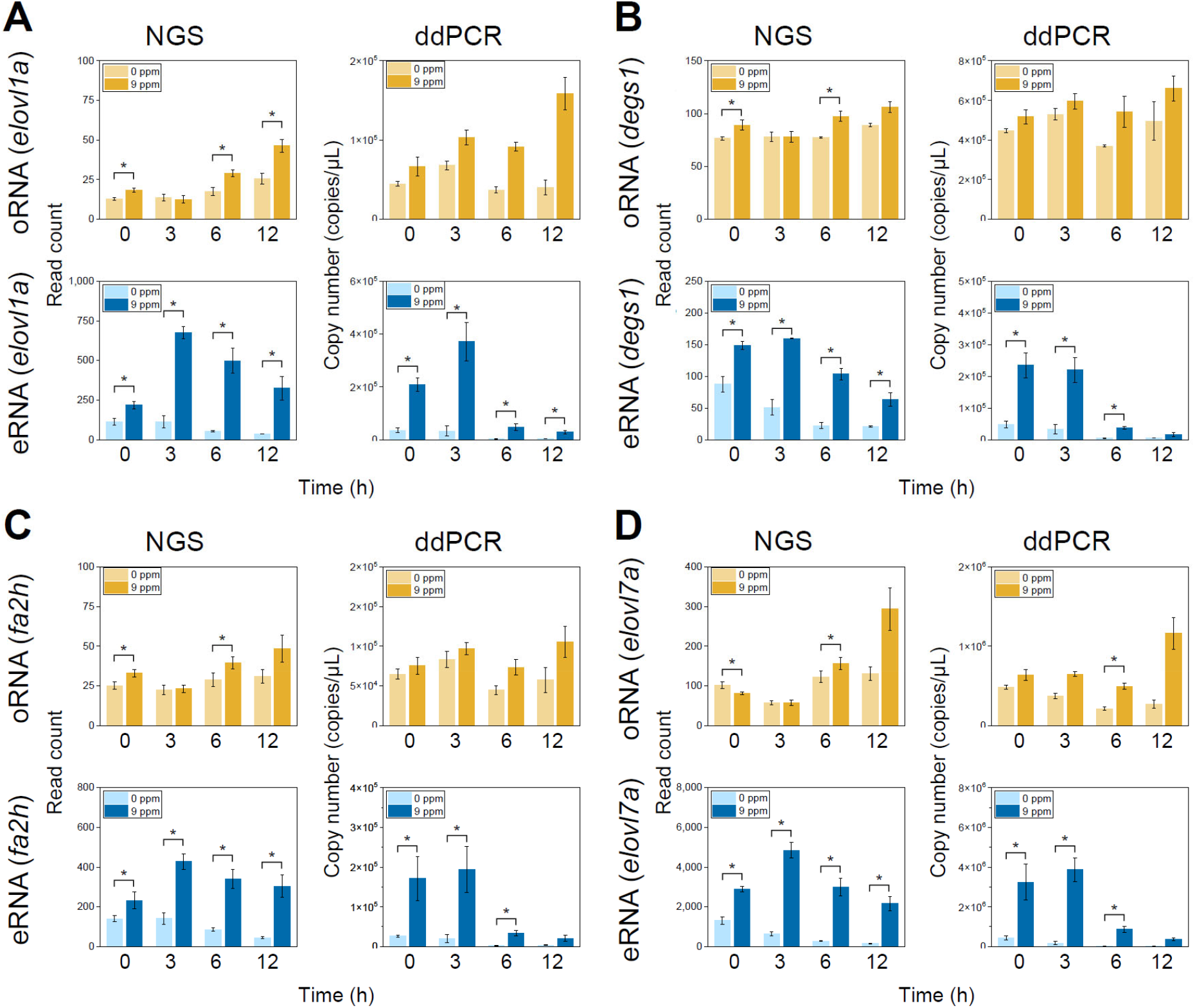
Differentially expressed gene identification and RT-ddPCR validation involving sphingolipid and ceramide synthesis. Read counts or copy numbers of (**A**) *elovl1a*, (**B**) *degs1*, (**C**) *fa2h*, and (**D**) *elovl7a* analyzed by NGS or ddPCR. Bar graphs showing averaged copy numbers of oRNA and eRNA in triplicate with (yellow or blue) or without LAS (light yellow or light blue). Error bars represent the standard deviation (SD). Differences between LAS-exposed and non-exposed groups were analyzed using paired-sample *t*-test. Asterisks indicate factors that are statistically significant after applying false discovery rate–adjusted *p* (*q*) values (\**q* < 0.1).

## 4. Discussion

Our eRNA-seq protocol, designed to assess optimal sampling time and mapping rates over time (Fig. S1), achieved mapping rates of 0.4–20.1% in treated samples (Table S5). By shortening the exposure period, we improved the mapping rates to 9.0–41.9% (Fig. 1E, Table S8), nearly 100 times higher than that reported in previous studies on *Daphnia pulex* (0.5%)^29^ and *O. latipes* (0.5%).^30^ This improvement was due to the fact that aquarium water was collected when eRNA degradation was minimal. Over time, total eRNA increased due to cell shedding and mucus production from LAS exposure (Fig. 1A), while bacterial degradation decreased eRNA quality (Fig. 1B), reducing the mapping rate after 24 h (Table S5). Notably, mapping rates were higher in the LAS-treated group, possibly due to increased RNA release as a result of enhanced RNA release in polluted environments. Consequently, our protocol facilitate crucial gene detection and detailed enrichment analyses, greatly improving the correlation between eRNA and oRNA read counts (R^2^: 0.5 ≤ 0.56) (Figs. 2B and S2) and the number of detected genes (Figs. 2A and S2) (1,110 (4%) ≤ 9,854 (38.2%)) compared to previous report,^30^ suggesting that eRNA collected within the initial 12 h provides high-resolution data.

Our protocol with high mapping rates was revealed that eRNA contains numerous intracellular genes, including nuclear ones (Figs. 2C and D). In this study, we detected 86.1% (22,216) of the 25,795 *O. latipes* genes in either oRNA or eRNA (Fig. 2A). Among 22,173 genes in oRNA, 9,811 (44.2%) were also found in eRNA. GO analysis revealed that oRNA genes predominantly encoded extracellular proteins, while eRNA mainly comprised intracellular genes, including those located in the nucleus (Figs. 2C and D). While Hiki et al. suggested challenges in identifying genes of nuclear origin from eRNA^30^, our findings indicate the detection of genes within the nucleus based on subcellular localization (Fig. 2C). This suggests that eRNA may originate from cells shed from feces and scales, similar to eDNA.^7–9,44–48^ Nuclear proteins are typically synthesized on ribosomes in the cytoplasm, differing from proteins produced on ribosomes in the rough endoplasmic reticulum.^49,50^ The resilience of nuclear proteins to external influences, such as LAS in this study, may contribute to their detectability in samples where RNA degradation is relatively less advanced. Furthermore, our analysis of gene expression in oRNA and eRNA, through the GO analysis on BP, showed distinct properties (Fig. S3, Table S9). These results suggest that eRNA has the capability to capture specific biological reactions that occur in individual organisms. The differences in BP detected may signify the variations in the organs from which the target cells are derived.

Enrichment analyses, such as GO and pathway analyses, require detecting a sufficient number of DEGs. We demonstrated that sampling aquarium water when eRNA degradation is minimal increases gene detection, allowing comprehensive chemical toxicity analysis through enrichment analysis. Our protocol identified more DEGs from eRNA (upregulated: 89–1,189, downregulated: 458–1,816) than those reported by Hiki et al. (upregulated: 196, downregulated: 115; Fig. S4).^30^ Interestingly, GO analysis on BP showed that functional terms associated with DEGs in oRNA changed over time, while those associated with eRNA remained consistent throughout the duration (Figs. 3A and B). It is likely that oRNA provides a snapshot of gene expression at a specific moment, whereas eRNA reflects cumulative gene expression over time, since eRNA was suggested to have the ability to assess the accumulation of time-varying gene expression.

Fish shows oxidative stress from LAS exposure due to ROS production.^32,33^ This, along with exposure to metals and surfactants, typically induces xenobiotic metabolism as well as inflammatory and immune responses in aquatic organisms.^51,52^ Our enrichment analysis of oRNA showed downregulation of xenobiotic metabolic processes (Fig. 4A) and upregulation of the glutathione metabolic process, indicating LAS-induced oxidative stress in *O. latipes*.^52,53^ Oxidative stress triggers sphingolipid and ceramide biosynthesis, which are associated with cell death, proliferation, and inflammation,^54–56^ and implicates EGFR in these processes.^57,58^ In this study, LAS exposure significantly activated sphingolipid and ceramide biosynthesis and upregulated the EGFR activity in oRNA. Autophagy, which is critical for disposing off damaged organelles, was also upregulated,^59–62^ while the MAPK cascade, a significant modulator of autophagy in response to oxidative stress,^63,64^ was downregulated (Fig. 4A and Table S12). The inhibition of the NF-κB signaling pathway, which is crucial for inflammation and immunity,^65^ along with cytokine-mediated signaling pathways and *IL-1β*, suggests that the LAS-induced oxidative stress induces the activation of immune-related signaling pathways (Fig. S5).^66^

Similar to oRNA, eRNA detected functions related to inflammation and immune responses, such as sphingolipid and ceramide biosynthesis and the EGFR signaling pathway. Additionally, eRNA exclusively detected functions, including protein glycosylation, sphingosine biosynthesis, cell–cell junction organization, and wound healing (Fig. 4B and Table S12). Protein glycosylation affects cell interactions and signaling,^67^ and disruptions in this phenomenon contribute to intestinal barrier dysfunction.^68^ The increased negative regulation of intrinsic apoptotic signaling aligns with apoptosis induced by sphingosine,^69^ which represents sphingolipid backbone. Wound healing processes^70,71^ were also upregulated, consistent with oRNA responses. Four out of five wound healing-related genes (*dsp*, *evpla*, *evplb*, and *ppl*) identified in oRNA (Fig. S6) were included in DEGs related to LAS effects (PC2 in PCA analysis; Fig. 5). The *ppl*, *evpla*, and *evplb* genes are expressed in gills and epithelial cells, while *dsp* is expressed in the intestinal epithelium.^72–76^ Given the continuous exposure of fishes to microorganisms, they develop mucus-associated lymphoid tissues in gills, intestines, and skin to protect against infection.^77^ Thus, genes detected in eRNA in aquarium water likely originate from these organs. Consequently, many functions related to inflammation were activated by LAS exposure and detected in eRNA, underscoring its utility as a non-invasive tool for monitoring cellular responses to chemical exposure in fish. Fold changes of genes were higher in eRNA than in oRNA, suggesting that eRNA is more sensitive than oRNA in detecting inflammatory responses (Figs. 6 and S6). Unexpectedly, autophagosome assembly was downregulated in eRNA, possibly due to gene degradation over time in the water. Thus, GO terms identified as downregulated in eRNA might not represent actual downregulation but rather degradation.

Therefore, caution is needed when interpreting eRNA-detected downregulation, as it is difficult to discern whether it is genuine or due to degradation, even with significant expression changes.

Focusing on upregulated genes, we observed a significant increase in sphingolipid and ceramide biosynthetic processes in response to LAS exposure in both oRNA and eRNA. Key enzymes essential for these processes, such as *elovl1a* and *elovl7a* (*elovl* family), *degs1* and *degs2* (*degs* family), *fa2h*, and *cersa* family, showed upregulation in both datasets (Fig. 6), suggesting potential production within the fish body.^78–81^ Notably, genes such as *elovl1a*, *degs1*, and *fa2h* were among the DEGs linked to the effects of LAS (PC2 in PCA analysis; Fig. 5) and are known to be expressed in skin and epithelial tissues in mammals.^79,82–86^ These results highlight the potential use of sphingolipid and ceramide biosynthetic genes as stress markers for assessing the effects of LAS exposure on *O. latipes*. These genes may have been detected as a result of cell shedding from body surface tissues or secretion into the water. If organisms release cells into the water in response to LAS exposure, eRNA from these cells could provide early insights into LAS effects. This suggests that detecting these genes in eRNA could offer earlier insights into LAS effects compared to oRNA, provided the following limitations are addressed. First, a sufficient amount of mRNA must be obtained from the real environment. Due to the presence of many RNA-degrading enzymes in natural settings, it is essential to ensure that the collected mRNA is both quantitatively and qualitatively adequate for analysis. Second, as LAS concentrations in real environments are much lower^87^ than those in this study, it is necessary to verify whether the expressions and properties of these gene in the eRNA can be detected at such concentrations. If these limitations can be overcome, eRNA-seq technology could become a valuable tool for the sensitive detection of stress in organisms in real environments.

## 5. Conclusion

eRNA-seq in chemical exposure experiments achieves a high mapping rate with well-timed sampling. This study significantly advances stress condition assessment in fish through eRNA gene expression analysis in aquarium water. Wound healing-related genes expressed in gills, gut, and skin cells were identified in both oRNA and eRNA datasets, demonstrating the non-invasive potential of eRNA-seq for understanding the stress status of aquatic organisms. While eDNA metabarcoding and traditional surveys have been valuable for species identification, they have not been able to determine stress status. eRNA-seq technology now addresses this gap, enabling the assessment of both species’ presence and stress status. These advances support biodiversity conservation aligned with the nature-positive framework and integrate fish health management in aquaculture, contributing to global food security.

## Author Contributions

**Kaede Miyata:** Conceptualization, Methodology, Investigation, Visualization, Project administration, Writing—original draft, Writing—review & editing. **Yasuaki Inoue:** Conceptualization, Methodology, Investigation, Visualization, Project administration, Writing— original draft, Writing—review & editing. **Masayuki Yamane:** Supervision, Writing—review & editing. **Hiroshi Honda:** Conceptualization, Supervision, Writing—review & editing.

## Acknowledgements

We thank Mr. Yuto Amano and Mr. Yuki Otsubo for their valuable discussions and assistance in setting up the analysis environment for this study. We thank Mrs. Tomoko Yokomatsu for her support in managing the Japanese medaka breeding system and maintaining the aquarium.

## Appendix A. Supplementary data

Supplementary data to this article can be found online at https://doi.org/xxx

## Abbreviations

eRNA, environmental RNA; eRNA-seq, eRNA-sequencing; LAS, linear alkylbenzene sulfonate; mRNA, messenger RNA; miRNA, micro RNA; oRNA, organismal RNA; PCA, principal component analysis.

## References

(1) Bongaarts, J. Human Population Growth and the Demographic Transition. Philosophical Transactions of the Royal Society B: Biological Sciences 2009, 364 (1532), 2985–2990. 10.1098/rstb.2009.0137.

(2) Dirzo, R.; Young, H. S.; Galetti, M.; Ceballos, G.; Isaac, N. J. B.; Collen, B. Defaunation in the Anthropocene. Science 2014, 345 (6195), 401–406. 10.1126/science.1251817.

(3) Collen, B.; Whitton, F.; Dyer, E. E.; Baillie, J. E. M.; Cumberlidge, N.; Darwall, W. R. T.; Pollock, C.; Richman, N. I.; Soulsby, A.-M.; Böhm, M. Global Patterns of Freshwater Species Diversity, Threat and Endemism. Global Ecol. Biogeogr. 2014, 23 (1), 40–51. 10.1111/geb.12096.

(4) Peng, B.; Sheng, X.; Wei, G. Does Environmental Protection Promote Economic Development? From the Perspective of Coupling Coordination between Environmental Protection and Economic Development. Environ. Sci. Pollut. Res. 2020, 27 (31), 39135–39148. 10.1007/s11356-020-09871-1.

(5) Everett, T.; Ishwaran, M.; Ansaloni, G. P.; Rubin, A. Economic Growth and the Environment. https://mpra.ub.uni-muenchen.de/23585/ (accessed 2024-03-29).

6. UN Environment Programme. Decision Adopted by the Conference of the Parties to the Convention on Biological Diversity 15/4. Kunming-Montreal Global Biodiversity Framework. 2022.

(7) Bohmann, K.; Evans, A.; Gilbert, M. T. P.; Carvalho, G. R.; Creer, S.; Knapp, M.; Yu, D. W.; de Bruyn, M. Environmental DNA for Wildlife Biology and Biodiversity Monitoring. Trends Ecol. Evol. 2014, 29 (6), 358–367. 10.1016/j.tree.2014.04.003.

(8) Deiner, K.; Bik, H. M.; Mächler, E.; Seymour, M.; Lacoursière-Roussel, A.; Altermatt, F.; Creer, S.; Bista, I.; Lodge, D. M.; de Vere, N.; Pfrender, M. E.; Bernatchez, L. Environmental DNA Metabarcoding: Transforming How We Survey Animal and Plant Communities. Mol. Ecol. 2017, 26 (21), 5872–5895. 10.1111/mec.14350.

(9) Ficetola, G. F.; Miaud, C.; Pompanon, F.; Taberlet, P. Species Detection Using Environmental DNA from Water Samples. Biol. Lett. 2008, 4 (4), 423–425. 10.1098/rsbl.2008.0118.

(10) TNFD Discussion Paper. A Landscape Assessment of Nature-Related Data and Analytics Availability, 2022. https://framework.tnfd.global/wp-content/uploads/2022/06/TNFD-Data-Discussion-Paper-Mar-2022.pdf.

(11) Kelly, R. P.; Port, J. A.; Yamahara, K. M.; Martone, R. G.; Lowell, N.; Thomsen, P. F.; Mach, M. E.; Bennett, M.; Prahler, E.; Caldwell, M. R.; Crowder, L. B. Environmental Monitoring. Harnessing DNA to Improve Environmental Management. Science 2014, 344 (6191), 1455–1456. 10.1126/science.1251156.

(12) Dougherty, M. M.; Larson, E. R.; Renshaw, M. A.; Gantz, C. A.; Egan, S. P.; Erickson, D. M.; Lodge, D. M. Environmental DNA (eDNA) Detects the Invasive Rusty Crayfish *Orconectes rusticus* at Low Abundances. J. Appl. Ecol. 2016, 53 (3), 722–732. 10.1111/1365-2664.12621.

13. A. Foote, A. D.; Thomsen, P. F.; Sveegaard, S.; Wahlberg, M.; Kielgast, J.; Kyhn, L. A.; Salling, B.; Galatius, A.; Orlando, L.; Gilbert, M. T. P. Investigating the Potential Use of Environmental DNA (eDNA) for Genetic Monitoring of Marine Mammals. PLoS One 2012, 7 (8), e41781. 10.1371/journal.pone.0041781.

(13) Fukumoto, S.; Ushimaru, A.; Minamoto, T. A Basin-Scale Application of Environmental DNA Assessment for Rare Endemic Species and Closely Related Exotic Species in Rivers: A Case Study of Giant Salamanders in Japan. J. Appl. Ecol. 2015, 52 (2), 358–365. 10.1111/1365-2664.12392.

(14) Port, J. A.; O’Donnell, J. L.; Romero-Maraccini, O. C.; Leary, P. R.; Litvin, S. Y.; Nickols, K. J.; Yamahara, K. M.; Kelly, R. P. Assessing Vertebrate Biodiversity in a Kelp Forest Ecosystem Using Environmental DNA. Mol. Ecol. 2016, 25 (2), 527–541. 10.1111/mec.13481.

(15) Thomsen, P. F.; Kielgast, J.; Iversen, L. L.; Wiuf, C.; Rasmussen, M.; Gilbert, M. T. P.; Orlando, L.; Willerslev, E. Monitoring Endangered Freshwater Biodiversity Using Environmental DNA. Mol. Ecol. 2012, 21 (11), 2565–2573. 10.1111/j.1365-294X.2011.05418.x.

(16) Thomsen, P. F.; Kielgast, J.; Iversen, L. L.; Møller, P. R.; Rasmussen, M.; Willerslev, E. Detection of a Diverse Marine Fish Fauna Using Environmental DNA from Seawater Samples. PLoS One 2012, 7 (8), e41732. 10.1371/journal.pone.0041732.

(17) Yamamoto, S.; Minami, K.; Fukaya, K.; Takahashi, K.; Sawada, H.; Murakami, H.; Tsuji, S.; Hashizume, H.; Kubonaga, S.; Horiuchi, T.; Hongo, M.; Nishida, J.; Okugawa, Y.; Fujiwara, A.; Fukuda, M.; Hidaka, S.; Suzuki, K. W.; Miya, M.; Araki, H.; Yamanaka, H.; Maruyama, A.; Miyashita, K.; Masuda, R.; Minamoto, T.; Kondoh, M. Environmental DNA as a “Snapshot” of Fish Distribution: A Case Study of Japanese Jack Mackerel in Maizuru Bay, Sea of Japan. PLoS One 2016, 11 (3), e0149786. 10.1371/journal.pone.0149786.

(18) Yamanaka, H.; Minamoto, T. The Use of Environmental DNA of Fishes as an Efficient Method of Determining Habitat Connectivity. Ecol. Indic. 2016, 62, 147–153. 10.1016/j.ecolind.2015.11.022.

(19) Cristescu, M. E.; Hebert, P. D. N. Uses and Misuses of Environmental DNA in Biodiversity Science and Conservation. Annu. Rev. Ecol. Evol. Syst. 2018, 49 (1), 209–230. 10.1146/annurev-ecolsys-110617-062306.

(20) Ficetola, G. F.; Taberlet, P.; Coissac, E. How to Limit False Positives in Environmental DNA and Metabarcoding? Mol. Ecol. Resour. 2016, 16 (3), 604–607. 10.1111/1755-0998.12508.

(21) Jo, T.; Murakami, H.; Masuda, R.; Sakata, M. K.; Yamamoto, S.; Minamoto, T. Rapid Degradation of Longer DNA Fragments Enables the Improved Estimation of Distribution and Biomass Using Environmental DNA. Mol. Ecol. Resour. 2017, 17 (6), e25–e33. 10.1111/1755-0998.12685.

(22) Alberts, B.; Johnson, A.; Lewis, J.; Morgan, D.; Raff, M.; Roberts, K.; Walter, P. Molecular Biology of the Cell (6th Edition); 2014.

(23) Miyata, K.; Inoue, Y.; Amano, Y.; Nishioka, T.; Nagaike, T.; Kawaguchi, T.; Morita, O.; Yamane, M.; Honda, H. Comparative Environmental RNA and DNA Metabarcoding Analysis of River Algae and Arthropods for Ecological Surveys and Water Quality Assessment. Sci. Rep. 2022, 12 (1), 19828. 10.1038/s41598-022-23888-1.

(24) Miyata, K.; Inoue, Y.; Amano, Y.; Nishioka, T.; Nagaike, T.; Kawaguchi, T.; Morita, O.; Yamane, M.; Honda, H. Comparative Environmental RNA and DNA Metabarcoding Analysis of River Algae and Arthropods for Ecological Surveys and Water Quality Assessment. Sci. Rep. 2022, 12 (1), 19828. 10.1038/s41598-022-23888-1.

(25) Inoue, Y.; Miyata, K.; Yamane, M.; Honda, H. Environmental Nucleic Acid Pollution: Characterization of Wastewater Generating False Positives in Molecular Ecological Surveys. ACS EST Water 2023, 3 (3), 756–764. 10.1021/acsestwater.2c00542.

(26) Cristescu, M. E. Can Environmental RNA Revolutionize Biodiversity Science? Trends in Ecology & Evolution 2019, 34 (8), 694–697. 10.1016/j.tree.2019.05.003.

(27) Ikert, H.; Lynch, M. D. J.; Doxey, A. C.; Giesy, J. P.; Servos, M. R.; Katzenback, B. A.; Craig, P. M. High Throughput Sequencing of MicroRNA in Rainbow Trout Plasma, Mucus, and Surrounding Water Following Acute Stress. Front. Physiol. 2021, 11, 588313. 10.3389/fphys.2020.588313.

(28) Hechler, R. M.; Yates, M. C.; Chain, F. J. J.; Cristescu, M. E. Environmental Transcriptomics under Heat Stress: Can Environmental RNA Reveal Changes in Gene Expression of Aquatic Organisms? bioRxiv November 18, 2022, p 2022.10.06.510878. 10.1101/2022.10.06.510878.

(29) Hiki, K.; Yamagishi, T.; Yamamoto, H. Environmental RNA as a Noninvasive Tool for Assessing Toxic Effects in Fish: A Proof-of-Concept Study Using Japanese Medaka Exposed to Pyrene. Environ. Sci. Technol. 2023, 57 (34), 12654–12662. 10.1021/acs.est.3c03737.

(30) Maruyama, A.; Nakamura, K.; Yamanaka, H.; Kondoh, M.; Minamoto, T. The Release Rate of Environmental DNA from Juvenile and Adult Fish. PLoS One 2014, 9 (12), e114639. 10.1371/journal.pone.0114639.

(31) Gouda, A. M. R.; Hagras, A. E.; Okbah, M. A.; El-Gammal, M. I. Influence of the Linear Alkylbenzene Sulfonate (LAS) on Hematological and Biochemical Parameters of Nile Tilapia, Oreochromis Niloticus. Saudi Journal of Biological Sciences 2022, 29 (2), 1006–1013. 10.1016/j.sjbs.2021.09.074.

(32) Shukla, A.; Trivedi, S. P. Anionic Surfactant, Linear Alkyl Benzene Sulphonate Induced Oxidative Stress and Hepatic Impairments in Fish Channa Punctatus. Proc. Zool. Soc. 2018, 71 (4), 382–389. 10.1007/s12595-017-0223-1.

(33) Ministry of the Environment, Japan. Public water area-Water quality measurement result. http://www.env.go.jp/water/suiiki/ (accessed 2024-02-06).

(34) Bolger, A. M.; Lohse, M.; Usadel, B. Trimmomatic: A Flexible Trimmer for Illumina Sequence Data. Bioinformatics 2014, 30 (15), 2114–2120. 10.1093/bioinformatics/btu170.

(35) Simon, A. FastQC A Quality Control Tool for High Throughput Sequence Data. Babraham Bioinformatics. https://www.bioinformatics.babraham.ac.uk/projects/fastqc/ (accessed 2023- 07-05).

(36) Li, H.; Durbin, R. Fast and Accurate Short Read Alignment with Burrows–Wheeler Transform. Bioinformatics 2009, 25 (14), 1754–1760. 10.1093/bioinformatics/btp324.

(37) Li, H.; Durbin, R. Fast and Accurate Long-Read Alignment with Burrows–Wheeler Transform. Bioinformatics 2010, 26 (5), 589–595. 10.1093/bioinformatics/btp698.

(38) Li, H.; Handsaker, B.; Wysoker, A.; Fennell, T.; Ruan, J.; Homer, N.; Marth, G.; Abecasis, G.; Durbin, R. The Sequence Alignment/Map Format and SAMtools. Bioinformatics 2009, 25 (16), 2078–2079. 10.1093/bioinformatics/btp352.

(39) Robinson, M. D.; Oshlack, A. A Scaling Normalization Method for Differential Expression Analysis of RNA-Seq Data. Genome Biol. 2010, 11 (3), R25. 10.1186/gb-2010-11-3-r25.

(40) Robinson, M. D.; McCarthy, D. J.; Smyth, G. K. edgeR: A Bioconductor Package for Differential Expression Analysis of Digital Gene Expression Data. Bioinformatics 2010, 26 (1), 139–140. 10.1093/bioinformatics/btp616.

(41) Chen, Y.; Lun, A. T. L.; Smyth, G. K. From Reads to Genes to Pathways: Differential Expression Analysis of RNA-Seq Experiments Using Rsubread and the edgeR Quasi-Likelihood Pipeline. F1000Res. 2016, 5, 1438. 10.12688/f1000research.8987.2.

(42) Huang, D. W.; Sherman, B. T.; Lempicki, R. A. Systematic and Integrative Analysis of Large Gene Lists Using DAVID Bioinformatics Resources. Nat. Protoc. 2009, 4 (1), 44–57. 10.1038/nprot.2008.211.

(43) Marshall, N. T.; Vanderploeg, H. A.; Chaganti, S. R. Environmental (e)RNA Advances the Reliability of eDNA by Predicting Its Age. Sci. Rep. 2021, 11, 2769. 10.1038/s41598-021-82205-4.

(44) Pochon, X.; Zaiko, A.; Fletcher, L. M.; Laroche, O.; Wood, S. A. Wanted Dead or Alive? Using Metabarcoding of Environmental DNA and RNA to Distinguish Living Assemblages for Biosecurity Applications. PLoS One 2017, 12 (11), e0187636. 10.1371/journal.pone.0187636.

(45) Tsuri, K.; Ikeda, S.; Hirohara, T.; Shimada, Y.; Minamoto, T.; Yamanaka, H. Messenger RNA Typing of Environmental RNA (eRNA): A Case Study on Zebrafish Tank Water with Perspectives for the Future Development of eRNA Analysis on Aquatic Vertebrates. Environ. DNA 2021, 3 (1), 14–21. 10.1002/edn3.169.

(46) von Ammon, U.; Wood, S. A.; Laroche, O.; Zaiko, A.; Lavery, S. D.; Inglis, G. J.; Pochon, X. Linking Environmental DNA and RNA for Improved Detection of the Marine Invasive Fanworm *Sabella spallanzanii*. Front. Mar. Sci. 2019, 6. 10.3389/fmars.2019.00621.

(47) Wood, S. A.; Biessy, L.; Latchford, J. L.; Zaiko, A.; von Ammon, U.; Audrezet, F.; Cristescu, M. E.; Pochon, X. Release and Degradation of Environmental DNA and RNA in a Marine System. Sci. Total Environ. 2020, 704, 135314. 10.1016/j.scitotenv.2019.135314.

(48) Kimura, M.; Morinaka, Y.; Imai, K.; Kose, S.; Horton, P.; Imamoto, N. Extensive Cargo Identification Reveals Distinct Biological Roles of the 12 Importin Pathways. eLife 6, e21184. 10.7554/eLife.21184.

(49) Alberts, B.; Johnson, A.; Lewis, J.; Raff, M.; Roberts, K.; Walter, P. From RNA to Protein. In Molecular Biology of the Cell. 4th edition; Garland Science, 2002.

(50) Braeuning, A.; Schwarz, M. Regulation of Expression of Drug-Metabolizing Enzymes by Oncogenic Signaling Pathways in Liver Tumors: A Review. Acta Pharm. Sin. B 2020, 10 (1), 113–122. 10.1016/j.apsb.2019.06.013.

(51) Silvestre, F. Signaling Pathways of Oxidative Stress in Aquatic Organisms Exposed to Xenobiotics. J. Exp. Zool. A Ecol. Integr. Physiol. 2020, 333 (6), 436–448. 10.1002/jez.2356.

(52) Lushchak, V. I. Environmentally Induced Oxidative Stress in Aquatic Animals. Aquat. Toxicol. 2011, 101 (1), 13–30. 10.1016/j.aquatox.2010.10.006.

(53) Maceyka, M.; Spiegel, S. Sphingolipid Metabolites in Inflammatory Disease. Nature 2014, 510 (7503), 58–67. 10.1038/nature13475.

(54) Fabri, J. H. T. M.; de Sá, N. P.; Malavazi, I.; Del Poeta, M. The Dynamics and Role of Sphingolipids in Eukaryotic Organisms upon Thermal Adaptation. Prog. Lipid Res. 2020, 80, 101063. 10.1016/j.plipres.2020.101063.

(55) Nikolova-Karakashian, M. N.; Reid, M. B. Sphingolipid Metabolism, Oxidant Signaling, and Contractile Function of Skeletal Muscle. Antioxid. Redox Signal. 2011, 15 (9), 2501–2517. 10.1089/ars.2011.3940.

(56) Zhao, L.; Liang, J.; Chen, F.; Tang, X.; Liao, L.; Liu, Q.; Luo, J.; Du, Z.; Li, Z.; Luo, W.; Yang, S.; Rahimnejad, S. High Carbohydrate Diet Induced Endoplasmic Reticulum Stress and Oxidative Stress, Promoted Inflammation and Apoptosis, Impaired Intestinal Barrier of Juvenile Largemouth Bass (Micropterus Salmoides). Fish Shellfish Immunol. 2021, 119, 308–317. 10.1016/j.fsi.2021.10.019.

(57) Nitta, M.; Kozono, D.; Kennedy, R.; Stommel, J.; Ng, K.; Zinn, P. O.; Kushwaha, D.; Kesari, S.; Furnari, F.; Hoadley, K. A.; Chin, L.; DePinho, R. A.; Cavenee, W. K.; D’Andrea, A.; Chen, C. C. Targeting EGFR Induced Oxidative Stress by PARP1 Inhibition in Glioblastoma Therapy. PLoS One 2010, 5 (5), e10767. 10.1371/journal.pone.0010767.

(58) Pesonen, M.; Vähäkangas, K. Autophagy in Exposure to Environmental Chemicals. Toxicol. Lett. 2019, 305, 1–9. 10.1016/j.toxlet.2019.01.007.

(59) Zhang, T.; Ma, C.; Zhang, Z.; Zhang, H.; Hu, H. NF-κB Signaling in Inflammation and Cancer. MedComm 2021, 2 (4), 618–653. 10.1002/mco2.104.

(60) Zhang, W.; Liu, H. T. MAPK Signal Pathways in the Regulation of Cell Proliferation in Mammalian Cells. Cell Res. 2002, 12 (1), 9–18. 10.1038/sj.cr.7290105.

(61) Debnath, J.; Baehrecke, E. H.; Kroemer, G. Does Autophagy Contribute To Cell Death? Autophagy 2005, 1 (2), 66–74. 10.4161/auto.1.2.1738.

(62) Luo, Y.; Zou, P.; Zou, J.; Wang, J.; Zhou, D.; Liu, L. Autophagy Regulates ROS-Induced Cellular Senescence via P21 in a P38 MAPKα Dependent Manner. Exp. Gerontol. 2011, 46 (11), 860–867. 10.1016/j.exger.2011.07.005.

(63) McClung, J. M.; Judge, A. R.; Powers, S. K.; Yan, Z. P38 MAPK Links Oxidative Stress to Autophagy-Related Gene Expression in Cachectic Muscle Wasting. Am. J. Physiol. Cell Physiol. 2010, 298 (3), C542–C549. 10.1152/ajpcell.00192.2009.

(64) Cui, Q.; Chen, F.-Y.; Chen, H.-Y.; Peng, H.; Wang, K.-J. Benzo[a]Pyrene (BaP) Exposure Generates Persistent Reactive Oxygen Species (ROS) to Inhibit the NF-κB Pathway in Medaka (*Oryzias melastigma*). Environ. Pollut. 2019, 251, 502–509. 10.1016/j.envpol.2019.04.063.

(65) Liu, T.; Zhang, L.; Joo, D.; Sun, S.-C. NF-κB Signaling in Inflammation. Signal Transduct. Target Ther. 2017, 2, 17023. 10.1038/sigtrans.2017.23.

(66) Groux-Degroote, S.; Cavdarli, S.; Uchimura, K.; Allain, F.; Delannoy, P. Chapter Four - Glycosylation Changes in Inflammatory Diseases. In Advances in Protein Chemistry and Structural Biology; Donev, R., Ed.; Inflammatory Disorders, Part A; Academic Press, 2020; Vol. 119, pp 111–156. 10.1016/bs.apcsb.2019.08.008.

(67) Lewis, C. V.; Taylor, W. R. Intestinal Barrier Dysfunction as a Therapeutic Target for Cardiovascular Disease. Am. J. Physiol. Heart Circ. Physiol. 2020, 319 (6), H1227–H1233. 10.1152/ajpheart.00612.2020.

(68) Ahn, E. H.; Lee, M. B.; Seo, D. J.; Lee, J.; Kim, Y.; Gupta, K. Sphingosine Induces Apoptosis and Down-Regulation of *MYCN* in PAX3–FOXO1-Positive Alveolar Rhabdomyosarcoma Cells Irrespective of *TP53* Mutation. Anticancer Res. 2018, 38 (1), 71–76. 10.21873/anticanres.12193.

(69) Christian, L. M.; Graham, J. E.; Padgett, D. A.; Glaser, R.; Kiecolt-Glaser, J. K. Stress and Wound Healing. Neuroimmunomodulation 2006, 13 (5–6), 337–346. 10.1159/000104862.

(70) Gouin, J.-P.; Kiecolt-Glaser, J. K. The Impact of Psychological Stress on Wound Healing: Methods and Mechanisms. Immunol. Allergy Clin. North Am. 2011, 31 (1), 81–93. 10.1016/j.iac.2010.09.010.

(71) Buckley, B. A.; Gracey, A. Y.; Somero, G. N. The Cellular Response to Heat Stress in the Goby *Gillichthys* Mirabilis: A cDNA Microarray and Protein-Level Analysis. J. Exp. Biol. 2006, 209 (14), 2660–2677. 10.1242/jeb.02292.

(72) Whitehead, A.; Zhang, S.; Roach, J. L.; Galvez, F. Common Functional Targets of Adaptive Micro- and Macro-Evolutionary Divergence in Killifish. Mol. Ecol. 2013, 22 (14), 3780–3796. 10.1111/mec.12316.

(73) Vij, S.; Purushothaman, K.; Sridatta, P. S. R.; Jerry, D. R. Transcriptomic Analysis of Gill and Kidney from Asian Seabass (*Lates* Calcarifer) Acclimated to Different Salinities Reveals Pathways Involved with Euryhalinity. Genes 2020, 11 (7), 733. 10.3390/genes11070733.

75. A. Kalinin, A. E.; Idler, W. W.; Marekov, L. N.; McPhie, P.; Bowers, B.; Steinert, P. M.; Steven, C. Co-Assembly of Envoplakin and Periplakin into Oligomers and Ca^2+^-Dependent Vesicle Binding: Implications for Cornified Cell Envelope Formation in Stratified Squamous Epithelia. J. Biol. Chem. 2004, 279 (21), 22773–22780. 10.1074/jbc.M313660200.

(74) Ali, M. F. Z.; Kameda, K.; Kondo, F.; Iwai, T.; Kurniawan, R. A.; Ohta, T.; Ido, A.; Takahashi, T.; Miura, C.; Miura, T. Effects of Dietary Silkrose of *Antheraea* Yamamai on Gene Expression Profiling and Disease Resistance to *Edwardsiella tarda* in Japanese Medaka (*Oryzias* Latipes). Fish Shellfish Immunol. 2021, 114, 207–217. 10.1016/j.fsi.2021.05.001.

(75) Salinas, I. The Mucosal Immune System of Teleost Fish. Biology 2015, 4 (3), 525–539. 10.3390/biology4030525.

(76) Sun, S.; Cao, X.; Gao, J. C24:0 Avoids Cold Exposure-Induced Oxidative Stress and Fatty Acid β-Oxidation Damage. iScience 2021, 24 (12), 103409. 10.1016/j.isci.2021.103409.

(77) Deák, F.; Anderson, R. E.; Fessler, J. L.; Sherry, D. M. Novel Cellular Functions of Very Long Chain-Fatty Acids: Insight From *ELOVL4* Mutations. Front. Cell Neurosci. 2019, 13, 428. 10.3389/fncel.2019.00428.

(78) Pant, D. C.; Dorboz, I.; Schluter, A.; Fourcade, S.; Launay, N.; Joya, J.; Aguilera-Albesa, S.; Yoldi, M. E.; Casasnovas, C.; Willis, M. J.; Ruiz, M.; Ville, D.; Lesca, G.; Siquier-Pernet, K.; Desguerre, I.; Yan, H.; Wang, J.; Burmeister, M.; Brady, L.; Tarnopolsky, M.; Cornet, C.; Rubbini, D.; Terriente, J.; James, K. N.; Musaev, D.; Zaki, M. S.; Patterson, M. C.; Lanpher, B. C.; Klee, E. W.; Pinto e Vairo, F.; Wohler, E.; Sobreira, N. L. de M.; Cohen, J. S.; Maroofian, R.; Galehdari, H.; Mazaheri, N.; Shariati, G.; Colleaux, L.; Rodriguez, D.; Gleeson, J. G.; Pujades, C.; Fatemi, A.; Boespflug-Tanguy, O.; Pujol, A. Loss of the Sphingolipid Desaturase DEGS1 Causes Hypomyelinating Leukodystrophy. J. Clin. Invest. 2019, 129 (3), 1240–1256. 10.1172/JCI123959.

(79) Wu, D.; Liu, Z.; Yu, P.; Huang, Y.; Cai, M.; Zhang, M.; Zhao, Y. Cold Stress Regulates Lipid Metabolism via AMPK Signalling in *Cherax* Quadricarinatus. J. Therm. Biol. 2020, 92, 102693. 10.1016/j.jtherbio.2020.102693.

(80) Tvrdik, P.; Westerberg, R.; Silve, S.; Asadi, A.; Jakobsson, A.; Cannon, B.; Loison, G.; Jacobsson, A. Role of a New Mammalian Gene Family in the Biosynthesis of Very Long Chain Fatty Acids and Sphingolipids. J. Cell Biol. 2000, 149 (3), 707–718. 10.1083/jcb.149.3.707.

(81) Sassa, T.; Ohno, Y.; Suzuki, S.; Nomura, T.; Nishioka, C.; Kashiwagi, T.; Hirayama, T.; Akiyama, M.; Taguchi, R.; Shimizu, H.; Itohara, S.; Kihara, A. Impaired Epidermal Permeability Barrier in Mice Lacking *Elovl1*, the Gene Responsible for Very-Long-Chain Fatty Acid Production. Mol. Cell. Biol. 2013, 33 (14), 2787–2796. 10.1128/MCB.00192-13.

(82) Ternes, P.; Franke, S.; Zähringer, U.; Sperling, P.; Heinz, E. Identification and Characterization of a Sphingolipid Δ4-Desaturase Family. J. Biol. Chem. 2002, 277 (28), 25512–25518. 10.1074/jbc.M202947200.

(83) Uchida, Y.; Hama, H.; Alderson, N. L.; Douangpanya, S.; Wang, Y.; Crumrine, D. A.; Elias, P. M.; Holleran, W. M. Fatty Acid 2-Hydroxylase, Encoded by *FA2H*, Accounts for Differentiation-Associated Increase in 2-OH Ceramides during Keratinocyte Differentiation. J. Biol. Chem. 2007, 282 (18), 13211–13219. 10.1074/jbc.M611562200.

(84) Eckhardt, M.; Yaghootfam, A.; Fewou, S. N.; Zöller, I.; Gieselmann, V. A Mammalian Fatty Acid Hydroxylase Responsible for the Formation of α-Hydroxylated Galactosylceramide in Myelin. Biochem. J. 2005, 388 (Pt 1), 245–254. 10.1042/BJ20041451.

(85) Shiode, S.; McDonough, K.; Belanger, S. E.; Carr, G. J. Probabilistic Environmental Risk Assessment for Linear Alkyl Benzene Sulfonate (LAS) in Japan Reduces Assessment Uncertainty. J. Water Environ. Technol. 2020, 18 (2), 80–94. 10.2965/jwet.19-016.

