## Supplementary Text, Figures, and Tables S1-8 for "Fish environmental RNA sequencing sensitively captures accumulative stress responses through short-term aquarium sampling"

for

**This PDF file includes:**

Supplementary Texts S1 to S4

Supplementary Information References

Figures S1 to S6

Tables S1 to S8

References (88 and 89)

CONTENTS

Text

Text S1. Estimation of the mapping rate at different sampling times to determine the sampling time

Text S2. Dynamics of eRNA released from *Oryzias latipes* in the aquarium to determine the sampling time

Text S3. Analysis of the effects of LAS concentration on *Oryzias latipes* to determine the LAS dosage

Text S4. LAS analysis

References

Figures

Fig. S1. Experimental design.

Fig. S2. Comparison of gene expression between oRNA and eRNA upon gene ontology analysis on cellular component ($p < 0.05$).

Fig. S3. Comparison of functional terms in the gene ontology analysis on biological process between oRNA and both oRNA and eRNA ($p < 0.05$).

Fig. S4. Number of DEGs between the LAS treatment and control groups.

Fig. S5. ddPCR-based evaluation of the copy numbers of *interleukin-1 beta* (*IL-1β*) that is involved in cytokine-mediated signaling pathway (GO:0019221)

Fig. S6. RT-ddPCR validation of the differentially expressed genes  involved in wound healing.

Tables

Table S1. Number of alive *Oryzias latipes* in the dose range experiments to estimate the appropriate exposure concentration.

Table S2. Body lengths and weights of *Oryzias latipes* samples.

Table S3. PCR primers employed for detecting *Oryzias latipes* and bacteria used in the sampling time finding experiments.

Table S4. ddPCR primers for amplifying DEGs related to the inflammatory response (wound healing, ceramide/sphingolipid biosynthesis process, and *IL-1β*) used in the main study.

Table S5. Summary of the mapping rate for finding the sampling time.

Table S6. Concentration of LAS at each time point.

Table S7. Number alive and conditions of water at the start and end points of the experiment.

Table S8. Summary of concentration, quality, and mapping rate.

Table S9. List of functional terms at 0 h without LAS in oRNA and eRNA in the gene ontology analysis on biological process in the same order as the bar plots shown in Figure S3. (XLSX: separate file)

Table S10. List of functional terms and -log $p$ value derived from up- and downregulated DEGs in oRNA and eRNA in the gene ontology analysis on biological process in the same order as in the balloon plots shown in Figures 3A and B. (XLSX: separate file)

Table S11. List of functional terms derived from DEGs in oRNA and eRNA in the gene ontology analysis on biological process in the same order as in the clustered heatmaps shown in Figure 3C. (XLSX: separate file)

Table S12. List of functional terms and component genes in the oRNA and eRNA in the gene ontology analysis on biological process in the same order as in the heatmaps shown in Figure 4. (XLSX: separate file)

### Supplementary Text

#### Text S1. Estimation of the mapping rate at different sampling times to determine the sampling time

The experimental methods are detailed as follows: The NIES_R strain of *Oryzias latipes* (*O. latipes*) utilized in our study was procured from the National Institute for Environmental Studies (Japan). This is bred in our laboratory for over two generations. The fish were maintained under consistent conditions, with a temperature of 24 ± 1 °C and a photoperiod of 16 h of light followed by 8 h of darkness. They were fed brine shrimp twice daily. Adult fish aged 3–4 months with similar body sizes were selected for the experiments.

Glass aquaria were prepared and filled with dechlorinated water, maintained at 24 ± 1 °C under a 16-hour light: 8-hour dark cycle. Fifteen fish were added to each aquarium, with negative controls prepared without fish. Water samples were collected at specified time points, filtered through a Sterivex cartridge, and processed accordingly. The main distinction lies in the duration of fish exposure (48 h in the previous part and 24 h in this part) and the corresponding sampling intervals.

At each sampling time point, triplicates of the glass aquarium and a negative control (containing dechlorinated water but no fish) were prepared. Water temperature, pH, and dissolved oxygen (DO) levels were measured using a digital thermometer (NETSUKEN Corp., Tokyo, Japan) and a pH/DO meter D-55 (HORIBA, Kyoto, Japan) before filtering. To minimize feces-related effects, no food was provided to the sample fish for 24 h prior to and during the experiment.

Total RNA extraction from water samples and fish was performed for both control and LAS treatment groups using ChargeSwitch® Total RNA Cell Kits (Thermo Fisher Scientific, Massachusetts, USA), followed by DNase I treatment to eliminate genomic DNA contamination. RNA was extracted from the entire fish body in each aquarium following the manufacturer's protocol, with three replicates prepared for each sampling point to reduce variability. Extraction of environmental RNA (eRNA) from water samples collected at each time point involved some modifications to the manufacturer's protocol. Briefly, 1 mL of lysis reagent was introduced into the Sterivex cartridge, followed by incubation at 60 °C for 1 h with gentle rotation. The resulting lysate was collected by centrifugation, and subsequent steps were performed according to the manufacturer's instructions. RNA concentration was quantified using a Qubit 2.0 Fluorometer (Life Technologies, CA, USA), while RNA quality was assessed using an Agilent 2200 TapeStation (Agilent Technologies, Inc., CA, USA).

After the experiment, the remaining fish (excluding those used in the study) were dissected and placed in 2 mL RNAlater (Life Technologies, Grand Island, NY, USA) overnight before being stored at −80 °C for future analysis. cDNA synthesis was conducted using the SMART-Seq v4 Ultra Low Input RNA Kit for Sequencing (TaKaRa Bio Inc., Shiga, Japan), with 3–10 ng of RNA from whole fish or water samples as input. Subsequently, cDNA was purified using AMPure XP beads (Beckman Coulter, Brea, CA, USA). A sequence library was generated using the Nextera XT DNA Library Prep Kit with Nextera XT Index Kit v2 Set A (Illumina, San Diego, CA, USA) according to the manufacturer's instructions, with 1 ng of cDNA from each sample used as input. All extracted RNA and cDNA samples were promptly stored at −80 °C for future use.

We generated a total of 30 sequencing libraries, comprising triplicate samples from three sampling time points, three treatment groups, and three quality controls. These libraries were constructed from both whole fish samples and water samples. Subsequently, the two sets of libraries underwent individual sequencing on a Nextseq 500/550 (Illumina, San Diego, CA, USA) platform using the NextSeq 500/550 High Output Kit v2.5 (150 cycles) (Illumina, San Diego, CA, USA), resulting in paired-end reads with a length of 150 base pairs (2 × 75 bp).

Raw reads were processed using Trimmomatic ver. 0.38 with the following parameters: ILLUMINACLIP: adapters.fa:2:30:10, LEADING:20, TRAILING:20, SLIDINGWINDOW:4:20, MINLEN:40.[35] Clean reads were obtained after filtering out low-quality reads and adaptors. Quality assessment of both raw and trimmed reads was conducted using FastQC ver 0.11.9 with default settings.[36] Clean reads were then aligned to the reference genome of *O. latipes* (accession number: GCF_002234675.1, accessed on February 28, 2023) using BWA ver. 0.7.17 with default parameters (Table S3).[37, 38] Read counts were quantified using SAMtools ver. 1.6 with default settings. Genes with read counts below 10 were filtered out using Microsoft Excel.[39] Normalization of read counts was performed using the trimmed mean of M values (TMM) method[40, 42] implemented in the edgeR ver. 3.16.5 package[41, 42] utilizing the calcNormFactors function. Mapping rate is calculated as mapped reads divided by total reads.

**Text S2. Dynamics of eRNA released from *Oryzias latipes* in the aquarium to determine the sampling time**

The setup for this experiment is similar to that described in the previous section about *O. latipes*. In both cases, *O. latipes* samples, obtained from the National Institute for Environmental Studies (Japan), were maintained under constant conditions and fed brine shrimp twice daily.

Glass aquaria were prepared and filled with dechlorinated water, maintained at 24 ± 1 °C under a 16-hour light: 8-hour dark cycle.

Prior to the commencement of the experiments, the glass aquaria (dimensions: 20 × 15 × 20 cm) were meticulously cleaned using detergent, a 1% chlorine bleach solution, and ethanol. Subsequently, they underwent multiple rinses with distilled water to eliminate any residual nucleic acid. Each aquarium was then filled with 3 L of dechlorinated water and randomly arranged on the water bath. The water temperature in the aquaria was maintained at 24 ± 1 °C using an open circuit chiller (Taitec Corp., Saitama, Japan), operating under a 16-h light: 8-h dark cycle, and continuously aerated using glass tubes and an air pump. Fifteen fishes were introduced into each aquarium and left undisturbed for up to 24 h. Additionally, negative controls were prepared, consisting of glass aquaria containing only dechlorinated water without any fish.

Water samples were collected from the entire aquaria at various time points, including pre-treatment, 0, 1, 3, 6, 12, and 24 h. After removing the fish, 3 L water samples were filtered through a Sterivex cartridge (0.45 µm; Millipore SVHV010RS; Merck, Tokyo, Japan) using a Masterflex peristaltic pump (Cole-Parmer, Vernon Hills, IL, USA).

Triplicates of the glass aquaria and negative controls were prepared at each sampling time point. Prior to filtration, the water temperature, pH, and DO levels were measured using a digital thermometer (NETSUKEN Corp., Tokyo, Japan) and a pH/DO meter D-55 (HORIBA, Kyoto, Japan). The fish were not fed for 24 h before the commencement of the experiment until its conclusion to control for the effect of feces.

Total DNA/RNA coextraction was carried out using the AllPrep DNA/RNA Mini Kit. Buffer RLT Plus (1 mL) was introduced into the Sterivex cartridge inlet. The cartridge underwent incubation with mild rotation at 60 °C for 60 min. The resulting lysis buffer was collected by centrifuging the Sterivex cartridge in a 50 mL tube at 8,000 g for 1 min, followed by the transfer of the supernatant to an AllPrep DNA spin column positioned in a 2 mL collection tube. Subsequent steps were executed according to the manufacturer's protocol, with an elution volume of 50 µL. Before cDNA synthesis, RNA samples were subjected to two consecutive DNase treatments using an rDNase set and a NucleoSpin RNA Clean-up XS kit (MACHEREY-NAGEL, Düren, Germany). Single-strand RNA was then reverse transcribed into cDNA using a PrimeScript II 1st Strand cDNA Synthesis Kit (TaKaRa Bio Inc., Shiga, Japan), following the manufacturer's protocol. DNA and RNA were simultaneously extracted from deionized water to evaluate cross contamination during sample filtration, DNA/RNA coextraction, and cDNA synthesis. All extracted RNA and cDNA samples were promptly stored at −80 °C.

To enhance the detection and identification of eRNA from *O. latipes*, a set of PCR primers was designed based on available reference sequences for the mitochondrial cytochrome b (*cytb*) gene.[88] The NIES_R strain utilized in this study comprises multiple subclades in the southern medaka, prompting the design of primers and probes targeting sequences common to each subclade identified through blast searches. The forward and reverse primers amplified a 210-base-pair (bp) fragment. This primer pair was employed for droplet digital PCR (ddPCR) along with a probe (see below). Sets of primers and TaqMan probes were procured from Invitrogen (Invitrogen, Carlsbad, CA, USA), and the primer sequences are provided in Table S3.

The cDNA was quantified using a Bio-Rad QX200 AutoDG Droplet Digital PCR system (Bio-Rad Laboratories, Hercules, CA, USA), which included negative filtration, negative coextraction, and negative cDNA synthesis controls. Prior to ddPCR analysis, cDNA was diluted at 1:10 to mitigate inhibition. PCR was conducted in a 22 µL reaction volume, comprising 11 µL of 2 × ddPCR Supermix for Probe, 1.1 μL of template cDNA, 2.0 μL of NIES_R forward and reverse primer pairs (10 μM), 0.6 μL of the NIES_R probe (Table S3), and 5.3 μL of sterile deionized water. Thermal cycling consisted of 95 °C for 5 min; 40 cycles of 95 °C for 30 s and 55 °C for 1 min; and a final step of 4 °C for 5 min followed by 95 °C for 5 min.

Similarly, the quantitation of bacteria over time in the aquarium experiment was carried out using ddPCR. PCR was conducted in a 22 μL reaction volume, comprising 11 μL of 2 × ddPCR Supermix for EvaGreen, 1.1 μL of template DNA, 0.3 μL of forward and reverse primer pairs 341F and R806 (10 μM) (Table S3),[89] and 9.3 μL of sterile deionized water. Thermal cycling conditions were 95 °C for 5 min; 50 cycles of 95 °C for 30 s and 55 °C for 2 min; followed by a final step of 4 °C for 5 min and 95 °C for 5 min. Each sample was quantified in triplicate, with negative controls (1.1 μL of sterile distilled water) and positive controls included in each plate. The positive control consisted of 1.1 μL of total DNA extracted from *O. latipes*. Data analysis was performed using Bio-Rad QuantaSoft software version 1.7.4 (Bio-Rad Laboratories). Samples with less than 10,000 accepted droplets were excluded from the analysis and re-measured until 10,000 accepted droplets were obtained. The total number of gene copies per reaction was calculated and converted to gene copies per 3 L of aquarium water sample. The bacterial growth curve data were fitted to a logistic model using OriginPro 2021b.

### Text S3. Analysis of the effects of LAS concentration on *Oryzias latipes* to determine the LAS dosage

The setup for this experiment is similar to that described in the previous section with respect to *O. latipes*. In both cases, *O. latipes* obtained from the National Institute for Environmental Studies (Japan) were maintained under constant conditions and fed brine shrimp twice daily.

Glass aquaria were prepared and filled with dechlorinated water, maintained at 24 ± 1 °C under a 16-h light: 8-h dark cycle.

Prior to the commencement of the experiments, the glass aquaria (dimensions: 20 × 15 × 20 cm) were meticulously cleaned using ethanol. Subsequently, they underwent multiple rinses with distilled water. The exposure concentrations included control and LAS-treated groups, and were set at 8, 9, and 10 mg/L. The effects of these LAS concentrations on *O. latipes* were observed. Each aquarium was then filled with 3 L of dechlorinated water and randomly arranged on the water bath. The water temperature in the aquaria was maintained at 24 ± 1 °C using an open circuit chiller (Taitec Corp., Saitama, Japan), operating under a 16-h light: 8-h dark cycle, and continuously aerated using glass tubes and an air pump. Fifteen fishes were introduced into each aquarium and left undisturbed for up to 12 h. At each time point, the number of alive *O. latipes* were counted.

**Text S4. LAS analysis**

LAS analysis was performed using liquid chromatography tandem mass spectrometry (LC/MS/MS; HPLC: 1200 Series, MS/MS: 6460 Triple Quad, Agilent Technologies, Santa Clara, CA, USA) at each sampling time point. C12-LAS-13C6 was added to each water sample as a surrogate standard before column separation. The solid-phase extraction cartridges (400 mg; Sep-Pak tC18 Plus, Waters, MA, USA) were conditioned with 10 mL MeOH, and then, 10 mL ultrapure water at 5 mL/min. Water samples (20 mL) were passed through the cartridge at 5 mL/min. The cartridge was then cleaned with 5 mL ultrapure water and dried for approximately 10 min under gentle vacuum. The target compounds were eluted with 6 mL MeOH. The eluate was evaporated under a gentle stream of nitrogen at 40 °C. The samples were reconstituted in 8 mL acetonitrile:water 40:60% (v/v) solution. Mobile phases consisted of 50 mM ammonium formate solution with 0.1% formic acid (mobile phase A). Acetonitrile (mobile phase B), and chromatographic separation was achieved using a Mightysil RP-18GP column (KANTO CHEMICAL, Japan; 2.0 × 150 mm, 3 μm) at a flow rate of 0.2 mL/min. The gradient (% mobile phase B) was as follows: initial: 40%, 3 min: 40%, 20 min: 65%, 10 min: 95%, and 6.0 min: 40%. The column oven was maintained at 40 °C. The sample injection volume was set at 5 μL. MS/MS was performed using electrospray ionization operated in the negative-ion mode. Analytes were acquired through multiple reaction monitoring experiments. The monitored precursor/fragment ions were m/z 297/183, 311/183, 325/183, 339/183, 353/183, and 331.2/176 for C10-LAS, C11-LAS, C12-LAS, C13-LAS, C14-LAS, and C12-LAS-13C6, respectively. Fragmentor voltages of 180 and 215 V were used to obtain the m/z 183 ion for C10-14 LAS and the m/z 176 ion for C12-LAS-13C6, respectively. The internal standard concentration was 6.25 μg/L. The determination of the LAS was performed using an absolute calibration curve method. The calibration curve was made using six calibration standards at concentrations of 0.005, 0.01, 0.02, 0.1, 0.2, and 0.5 mg/L. Calibration curves were constructed using analyte/internal standard peak area ratio versus analyte concentration. For all the compounds, the values of coefficient of determination ($R^2$) of the calibration curves were 0.9992–0.9995. The limit of quantification of the LAS was the lowest concentration of the standard solution (0.005 mg/L) within the range of the confirmed calibration. Therefore, considering pretreatment, the limit of quantification of each component and the total LAS (C10-C14 LAS) were 0.002 and 0.01 mg/L, respectively.

### Supplementary Information References

(88) Takehana, Y.; Nagai, N.; Matsuda, M.; Tsuchiya, K.; Sakaizumi, M. Geographic Variation and Diversity of the Cytochrome b Gene in Japanese Wild Populations of Medaka, *Oryzias Latipes*. *Zoolog.* Sci. **2003**, 20 (10), 1279–1291. https://doi.org/10.2108/zsj.20.1279.

(89) Takahashi, S.; Tomita, J.; Nishioka, K.; Hisada, T.; Nishijima, M. Development of a Prokaryotic Universal Primer for Simultaneous Analysis of Bacteria and Archaea Using Next-Generation Sequencing. PLoS One 2014, *9* (8), e105592. https://doi.org/10.1371/journal.pone.0105592.

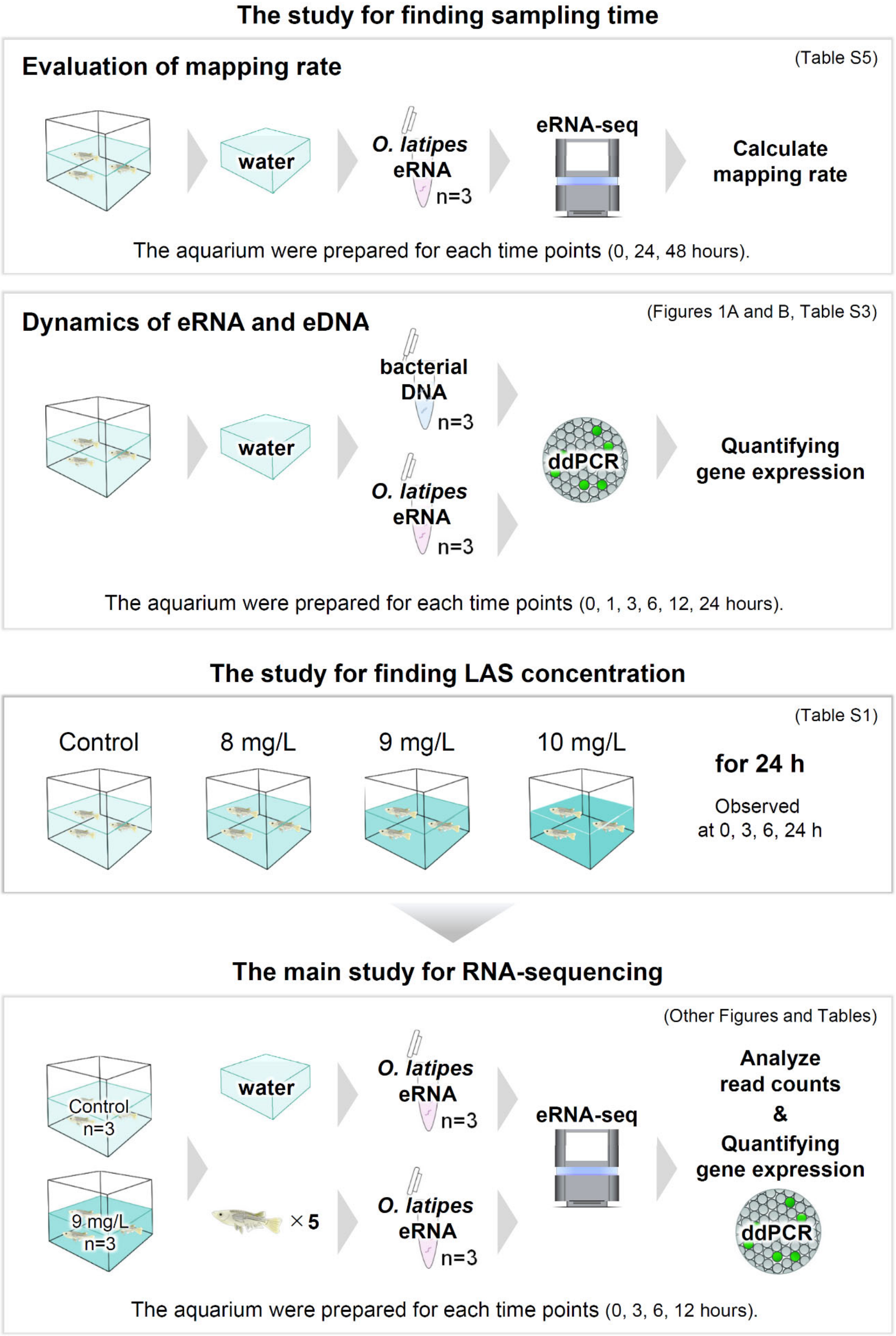
The study for finding sampling time
Evaluation of mapping rate
(Table S5)
water
O. latipes eRNA
n=3
eRNA-seq
Calculate mapping rate
The aquarium were prepared for each time points (0, 24, 48 hours).
Dynamics of eRNA and eDNA
(Figures 1A and B, Table S3)
water
bacterial DNA
n=3
O. latipes eRNA
n=3
ddPCR
Quantifying gene expression
The aquarium were prepared for each time points (0, 1, 3, 6, 12, 24 hours).
The study for finding LAS concentration
(Table S1)
Control
8 mg/L
9 mg/L
10 mg/L
for 24 h
Observed at 0, 3, 6, 24 h
The main study for RNA-sequencing
(Other Figures and Tables)
Control n=3
9 mg/L n=3
water
×5
O. latipes eRNA
n=3
O. latipes eRNA
n=3
eRNA-seq
Analyze read counts & Quantifying gene expression
ddPCR
The aquarium were prepared for each time points (0, 3, 6, 12 hours).

**Fig. S1. Experimental design.** Experiments were performed to detect the appropriate sampling time for further experiments. The quantity of bacterial DNA and *O. latipes* eRNA were measured using ddPCR in the water samples. The main experiments for RNA-seq were performed in triplicate at each time point (0, 3, 6, and 12 h after LAS exposure). Whole bodies of five pre- and post-stress fishes each were sequenced from each experimental replicate. One pre- and post-stress water sample each were sequenced from each experimental replicate.

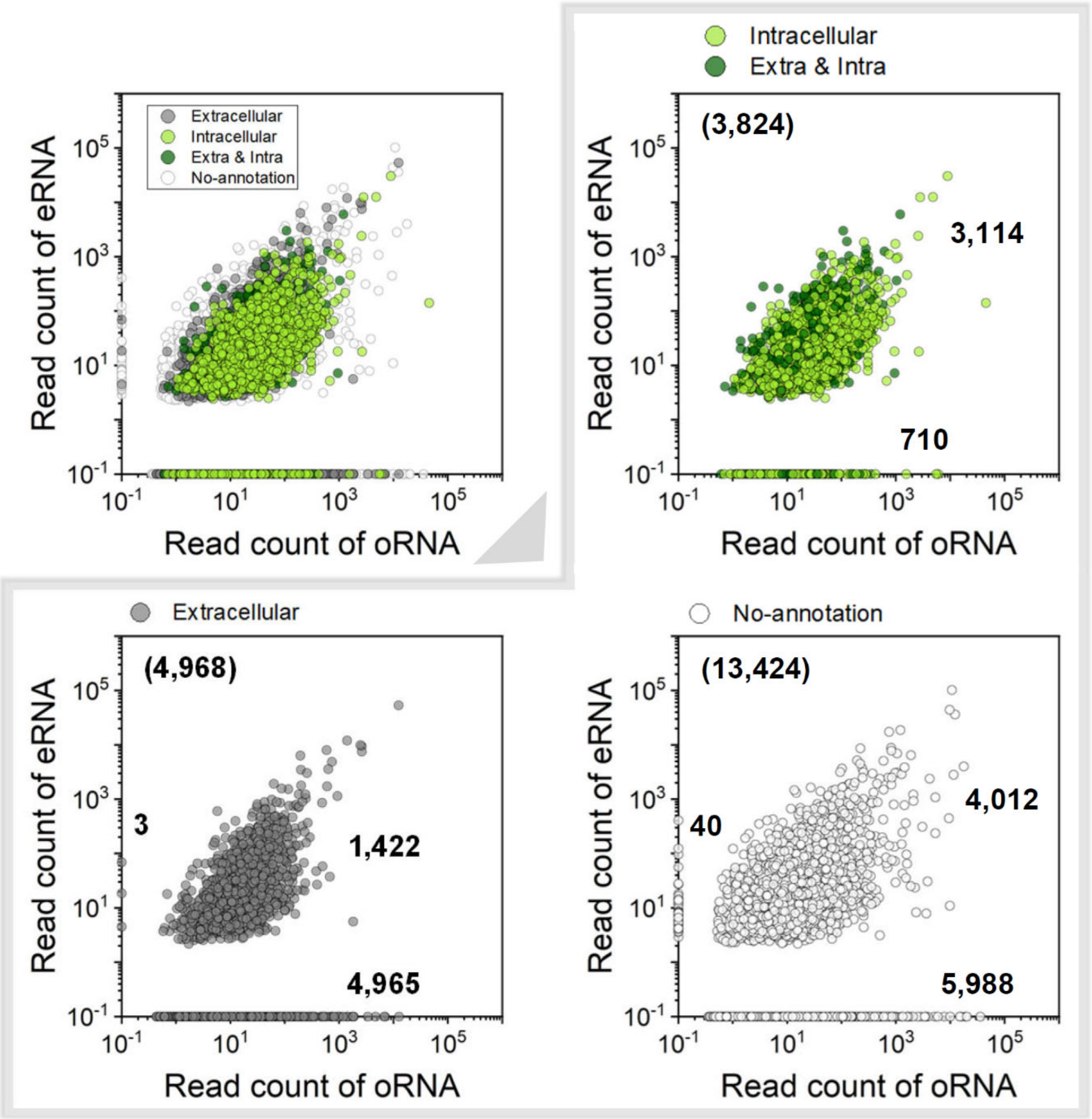


**Fig. S2. Comparison of gene expression between oRNA and eRNA upon gene ontology analysis on cellular component ($p < 0.05$).** Scatter plot showing read counts of all genes between oRNA and eRNA. Genes are categorized as extracellular (gray dots), intracellular (light green dots), both (green dots), and no-annotation (white dots). The numbers in the scatter plot indicate the number of genes at each location.

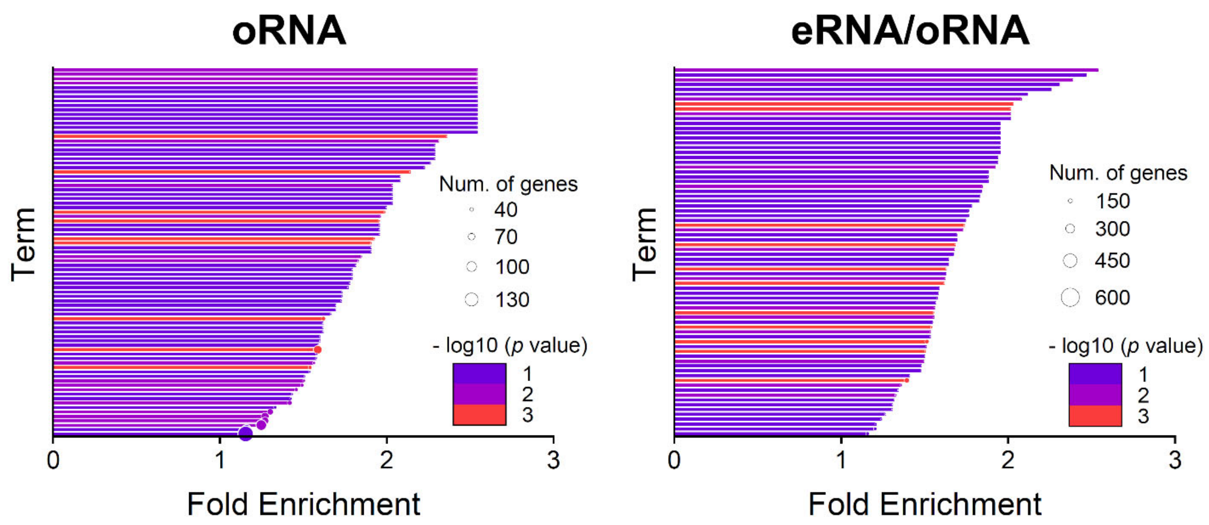


**Fig. S3. Comparison of functional terms in the gene ontology analysis on biological process between oRNA and both oRNA and eRNA ($p$ < 0.05).** Bar graph showing fold enrichment scores in each functional term. Color represents −log $p$ value of each term. Bubble size represents number of genes.

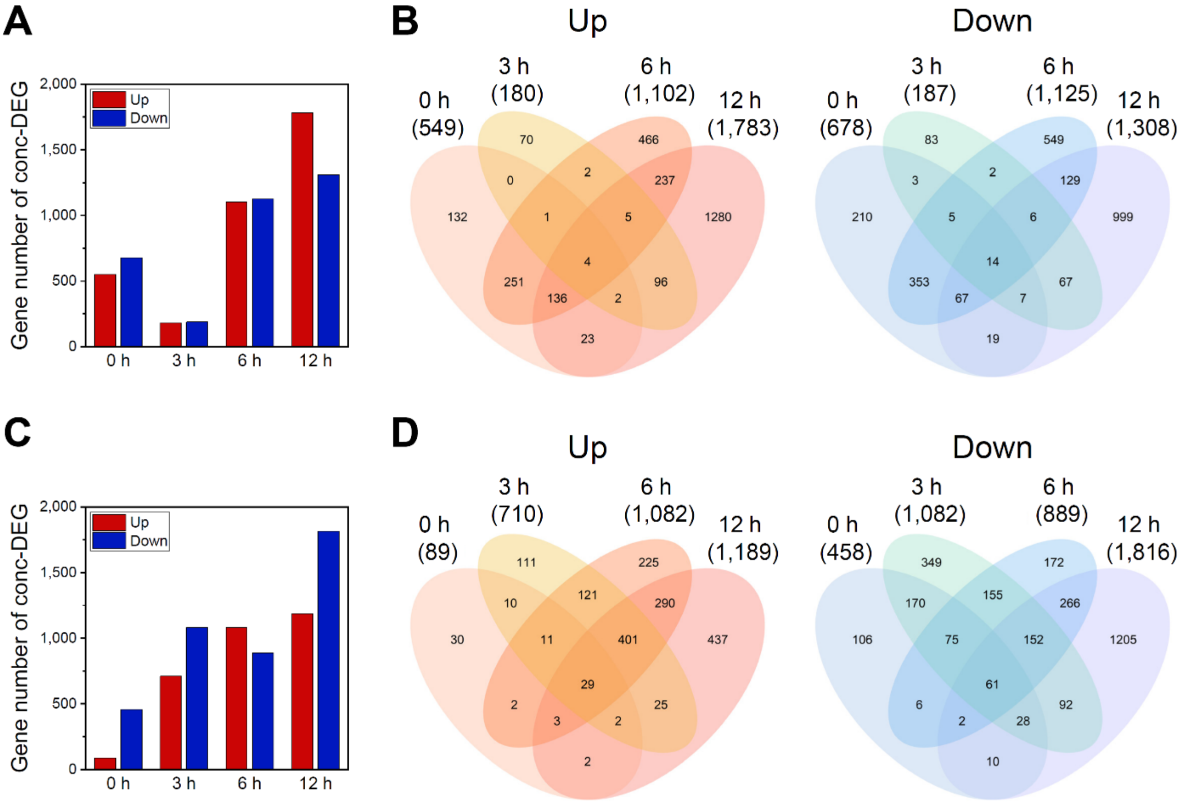


**Fig. S4. Number of DEGs between the LAS treatment and control groups.** (**A and C**) Bar graph showing the number of DEGs detected from (**A**) oRNA or (**C**) eRNA over time. Red bar: upregulated; Blue bar: downregulated. (**B and D**) Venn diagrams of the number of intersections of DEGs in (**B**) oRNA or (**D**) eRNA between each time point in up- (left) or downregulated genes (right).

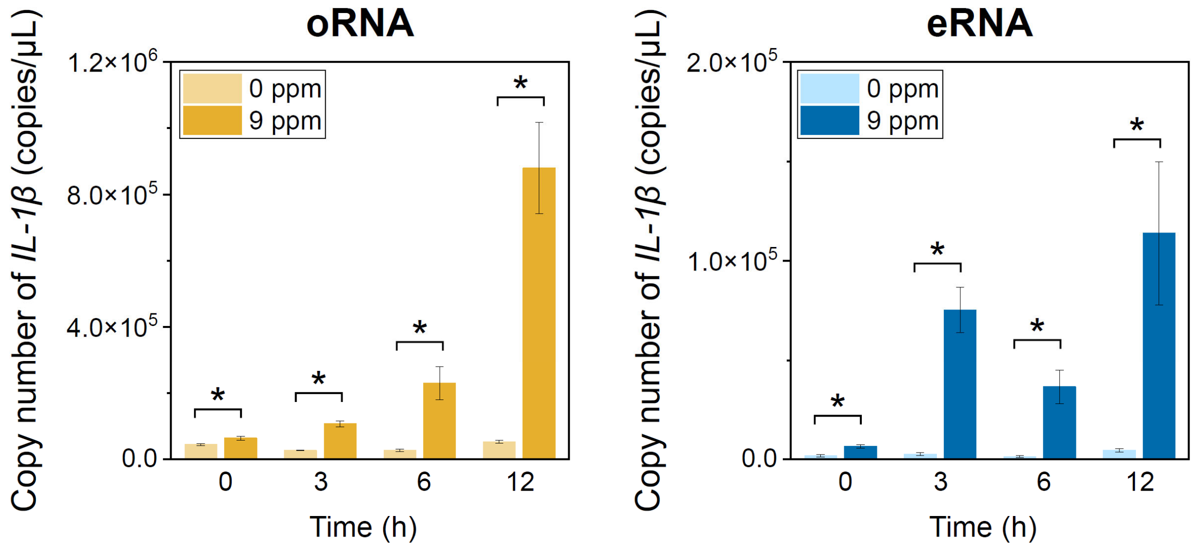


**Fig. S5. ddPCR-based evaluation of the copy numbers of *interleukin-1 beta* (*IL-1β*) that is involved in cytokine-mediated signaling pathway (GO:0019221) .** Bar graphs showing averaged copy numbers of oRNA and eRNA in triplicate with (yellow or blue) or without LAS (light yellow or light blue). Error bars show standard deviations (SD). Differences between with or without LAS exposure quantities were analyzed using paired-sample *t*-test. Asterisks indicate factors that are statistically significant after applying FDR-adjusted *p* (*q*) values (*$q < 0.1$).

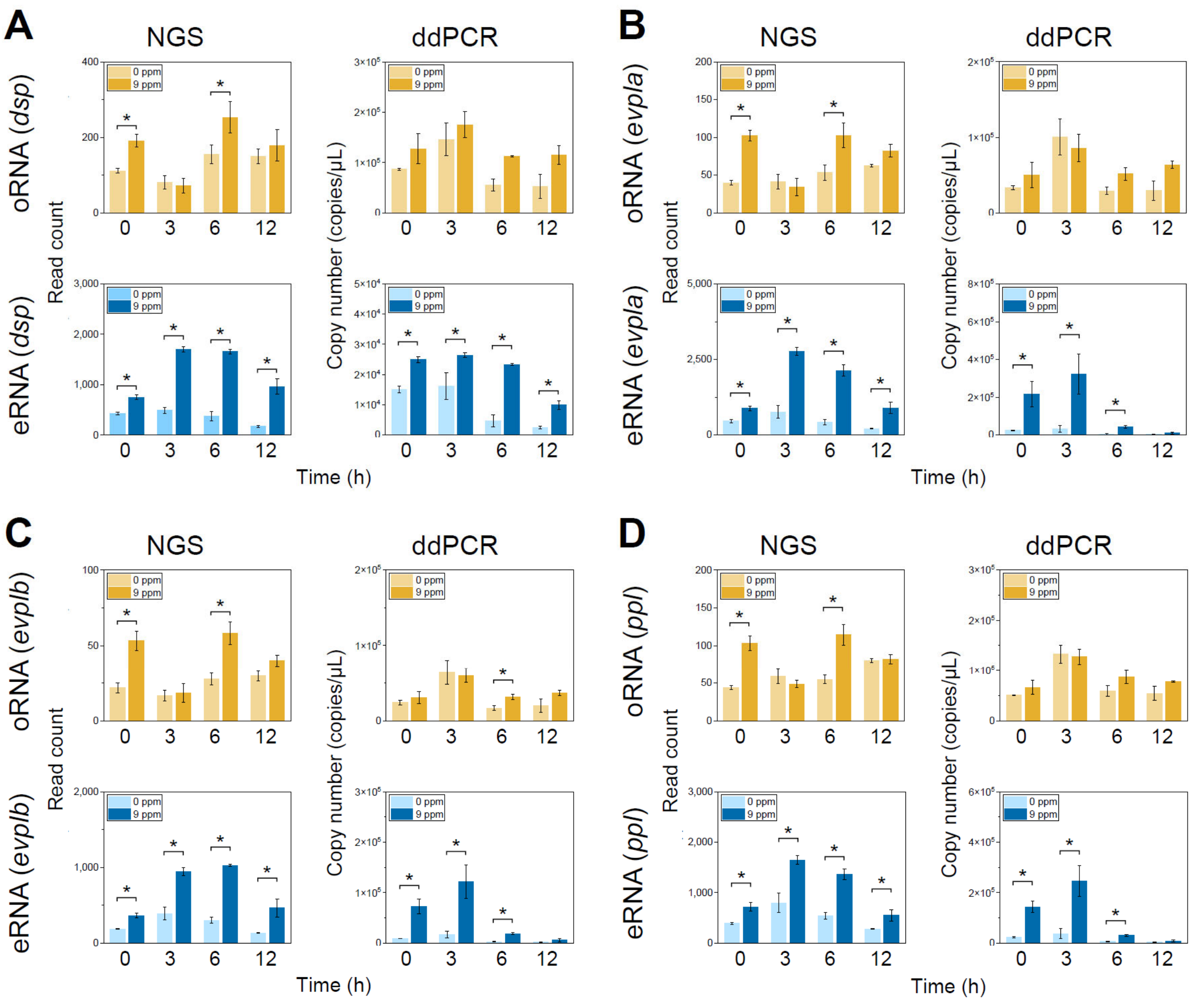


**Fig. S6. RT-ddPCR validation of the differentially expressed genes involved in wound healing.** Read counts or copy numbers of (**A**) *dsp*, (**B**) *evpla*, (**C**) *evplb*, and (**D**) *ppl* by NGS or ddPCR. Bar graphs showing averaged copy numbers of oRNA and eRNA in triplicate with (yellow or blue) or without LAS (light yellow or light blue). Error bars showing standard deviation (SD). Differences between with or without LAS exposure quantities were analyzed using paired-sample *t*-test. Asterisks indicate factors that are statistically significant after applying FDR-adjusted $p$ ($q$) values (*$q < 0.1$).

**Table S1. Number of alive *Oryzias latipes* in the dose range experiments to estimate the appropriate exposure concentration.** The input number of *O. latipes* was 15 in 3 L/aquarium.

| **LAS concentration (mg/L)** | **3 h** | **6 h** | **12 h** |
|---|---|---|---|
| 0 | 15 | 15 | 15 |
| 8 | 15 | 15 | 15 |
| 9 | 15 | 15 | 11 |
| 10 | 15 | 15 | 5 |

**Table S2. Body lengths and weights of *Oryzias latipes* samples.**

| Sample no. | Body length (cm) | Body weight (g) |
|---|---|---|
| 1 | 2.19 | 0.13 |
| 2 | 2.47 | 0.17 |
| 3 | 2.22 | 0.17 |
| 4 | 2.10 | 0.13 |
| 5 | 2.44 | 0.19 |
| 6 | 2.44 | 0.20 |
| 7 | 2.29 | 0.14 |
| 8 | 2.23 | 0.14 |
| 9 | 2.21 | 0.17 |
| 10 | 2.24 | 0.14 |
| Ave. | 2.28 | 0.16 |

**Table S3. PCR primers employed for detecting *Oryzias latipes* and bacteria used in the sampling time finding experiments.**

| **Species (target gene)** | **Primer name** | **Sequence (5′ to 3′)** | **Fluorophore /Quencher** | **Annealing temperature (°C)** |
|---|---|---|---|---|
| *Oryzias latipes (cytochrome b)* | NIES_R F | TTTGCCTACGCCATTCTACG | N/A | 55 |
| | NIES_R R | GGCTTCGTTGTTTAGAGGTGTG | N/A | |
| | probe | TTAGCCTCTATTCTAGTACTATTC | FAM/QSY | |
| Bacteria (*16S rRNA*) | 341F | CCTACGGGAGGCAGCAG | N/A | 55 |
| | R806 | GGACTACHVGGGTWTCTAAT | N/A | |

F; Forward, R; Reverse

**Table S4. ddPCR primers for amplifying DEGs related to the inflammatory response (wound healing, ceramide/sphingolipid biosynthesis process, and *IL-1β*) used in the main study.**

| Gene ID | Gene symbol | Primer | Sequence (5′→3′) | Annealing temperature (°C) |
|---|---|---|---|---|
| 101157810 | *elovl1a* | F | TCAAGGATTGATTCCAGGGT | 56 |
| | | R | TCGAATCATCCTGAGAGCCT | |
| 101175288 | *degs1* | F | AGGATATCCTGGCCAAACAC | 56 |
| | | R | AAGGCAGTGTTGTGGGAGAT | |
| 101160316 | *fa2h* | F | CTGGACAAGGATTTGGTCGA | 56 |
| | | R | GACCCAGTACCAGGAGGTTT | |
| 101167544 | *elovl7a* | F | ATGCAGATTCCAGGACAGAG | 56 |
| | | R | TTGCTGCCATCCTCATAGCT | |
| 101165763 | *dsp* | F | GACAAGAACATCGACCAGTA | 56 |
| | | R | CGTTCTTTGTCTTTCTCCAA | |
| 101167172 | *evpla* | F | AACTGACAACCCTGAAGAGG | 57 |
| | | R | GTCTCTCTGAGCCTGTATGG | |
| 101155032 | *evplb* | F | GCGAGTAGAACACCTTCTGT | 55 |
| | | R | GTGGAGATTCTTCACGTCTC | |
| 101174086 | *ppl* | F | CCAACAGGCTGTGAAGGACT | 58 |
| | | R | GACCTCGGGTTGCTGTTTTA | |
| 101169209 | *LOC101169209* (*IL-1β*) | F | GCAGAACCGGGGAGTATCAG | 59 |
| | | R | CTGTTGTCCTCCACCTCCTC | |

F; Forward, R; Reverse

**Table S5. Summary of the mapping rate for finding the sampling time.**

| Time | LAS (mg/L) | oRNA | | eRNA | |
|---|---|---|---|---|---|
| | | Raw reads (M) | Mapping rate (%) | Raw reads (M) | Mapping rate (%) |
| 10 min | 0 | 36.0 ± 1.19 | 92.9 ± 0.22 | 37.3 ± 2.20 | 11.4 ± 0.57 |
| 24 h | 0 | 38.9 ± 1.59 | 92.5 ± 1.53 | 33.5 ± 1.51 | 1.0 ± 0.19 |
| 48 h | 0 | 36.4 ± 2.70 | 92.2 ± 0.76 | 36.9 ± 3.18 | 0.8 ± 0.02 |
| 10 min | 4 | 33.4 ± 0.65 | 93.3 ± 1.22 | 37.6 ± 3.76 | 12.3 ± 0.41 |
| 24 h | 4 | 36.5 ± 1.63 | 93.2 ± 0.84 | 33.2 ± 2.29 | 0.4 ± 0.06 |
| 48 h | 4 | 32.6 ± 0.99 | 91.6 ± 1.33 | 31.0 ± 0.52 | 0.6 ± 0.14 |
| 10 min | 8 | 32.7 ± 2.19 | 92.7 ± 1.60 | 36.1 ± 0.17 | 20.1 ± 0.37 |
| 24 h | 8 | 35.5 ± 4.55 | 91.8 ± 0.25 | 40.2 ± 5.81 | 0.5 ± 0.03 |
| 48 h | 8 | 31.6 ± 2.38 | 91.2 ± 0.24 | 26.5 ± 2.03 | 0.6 ± 0.05 |

Average ± standard deviation (n = 1–3).

**Table S6. Concentration of LAS at each time point.** [a]Solvent control (Sol. con.) with no-fish at 10 min. Other samples collected from the aquarium with fish. LOD: Limit of detection.

| | LAS (mg/L) | C10-LAS | C11-LAS | C12-LAS | C13-LAS | C14-LAS | Total |
|---|---|---|---|---|---|---|---|
| Sol. con.[a] | 0 | < 0.002 | < 0.002 | < 0.002 | < 0.002 | < 0.002 | < 0.01 |
| 10 min | | < 0.002 | < 0.002 | < 0.002 | < 0.002 | < 0.002 | < 0.01 |
| 3 h | | < 0.002 | < 0.002 | < 0.002 | < 0.002 | < 0.002 | < 0.01 |
| 6 h | | < 0.002 | < 0.002 | < 0.002 | < 0.002 | < 0.002 | < 0.01 |
| 12 h | | < 0.002 | < 0.002 | < 0.002 | < 0.002 | < 0.002 | < 0.01 |
| LOD | | 0.002 | 0.002 | 0.002 | 0.002 | 0.002 | 0.01 |
| Sol. con.[a] | 9 | 0.87 | 2.72 | 2.77 | 2.11 | < 0.01 | 8.47 |
| 10 min | | 0.83 | 2.64 | 2.65 | 1.90 | < 0.01 | 8.02 |
| 3 h | | 0.79 | 2.51 | 2.51 | 1.76 | < 0.01 | 7.57 |
| 6 h | | 0.85 | 2.73 | 2.88 | 2.30 | < 0.01 | 8.76 |
| 12 h | | 0.85 | 2.66 | 2.69 | 1.93 | < 0.01 | 8.13 |
| LOD | | 0.01 | 0.01 | 0.01 | 0.01 | 0.01 | 0.05 |

**Table S7. Number alive and conditions of water at the start and end points of the experiment.**

| Time | LAS (mg/L) | n | Start | | | | End | | | |
|---|---|---|---|---|---|---|---|---|---|---|
| | | | Num. alive | Temp. (°C) | DO | pH | Num. alive | Temp. (°C) | DO | pH |
| 10 min | 0 | n = 1 | 0 | 24.3 | 8.57 | 7.01 | 0 | - | - | - |
| | | n = 2 | 0 | - | - | - | 0 | - | - | - |
| | | n = 3 | 0 | - | - | - | 0 | - | - | - |
| | 9 | n = 1 | 0 | 23.8 | 7.87 | 6.91 | 0 | - | - | - |
| | | n = 2 | 0 | - | - | - | 0 | - | - | - |
| | | n = 3 | 0 | - | - | - | 0 | - | - | - |
| 10 min | 0 | n = 1 | 15 | - | - | - | 15 | 24.8 | 8.55 | 7.28 |
| | | n = 2 | 15 | - | - | - | 15 | - | - | - |
| | | n = 3 | 15 | - | - | - | 15 | - | - | - |
| 3 h | 0 | n = 1 | 15 | - | - | - | 15 | 23.9 | 8.37 | 7.53 |
| | | n = 2 | 15 | - | - | - | 15 | - | - | - |
| | | n = 3 | 15 | - | - | - | 15 | - | - | - |
| 6 h | 0 | n = 1 | 15 | - | - | - | 15 | 23.9 | 7.94 | 7.47 |
| | | n = 2 | 15 | - | - | - | 15 | - | - | - |
| | | n = 3 | 15 | - | - | - | 15 | - | - | - |
| 12 h | 0 | n = 1 | 15 | - | - | - | 15 | 23.9 | 7.88 | 7.65 |
| | | n = 2 | 15 | - | - | - | 15 | - | - | - |
| | | n = 3 | 15 | - | - | - | 15 | - | - | - |
| 10 min | 9 | n = 1 | 15 | - | - | - | 15 | 24.3 | 8.67 | 7.31 |
| | | n = 2 | 15 | - | - | - | 15 | - | - | - |
| | | n = 3 | 15 | - | - | - | 15 | - | - | - |
| 3 h | 9 | n = 1 | 15 | - | - | - | 15 | 23.9 | 8.12 | 7.15 |
| | | n = 2 | 15 | - | - | - | 15 | - | - | - |
| | | n = 3 | 15 | - | - | - | 15 | - | - | - |
| 6 h | 9 | n = 1 | 15 | - | - | - | 15 | 23.9 | 7.70 | 6.93 |
| | | n = 2 | 15 | - | - | - | 15 | - | - | - |
| | | n = 3 | 15 | - | - | - | 15 | - | - | - |
| 12 h | 9 | n = 1 | 15 | - | - | - | 15 | 24.1 | 8.01 | 7.68 |
| | | n = 2 | 15 | - | - | - | 14 | - | - | - |
| | | n = 3 | 15 | - | - | - | 14 | - | - | - |

Num.: Number. Temp.: Temperature.

**Table S8. Summary of concentration, quality, and mapping rate of oRNA/eRNA.** The concentration and quality (RNA Integrity Number equivalent [RIN$^{e}$]) were measured using Qubit 4.0 Fluorometer and Agilent 2200 TapeStation, respectively.

| Time | LAS (mg/L) | oRNA | | | | eRNA | | | |
|---|---|---|---|---|---|---|---|---|---|
| | | Conc. (ng/µL) | RIN$^{e}$ | Raw reads(M) | Mapping rate (%) | Conc. (ng/µL) | RIN$^{e}$ | Raw reads (M) | Mapping rate (%) |
| Con.[*] | 0 or 9 | - | - | 0.010 ± 0.003 | 90.0 ± 7.89 | Low | 7.7 or N.D.[†] | 0.025 ± 0.002 | 94.6 ± 1.27 |
| 10 m | 0 | 2,787 ± 24.9 | 8.50 ± 0.08 | 39.8 ± 2.04 | 91.2 ± 0.36 | Low | 6.20 ± 0.08 | 40.5 ± 0.99 | 24.1 ± 1.41 |
| 3 h | | 2,627 ± 83.8 | 8.60 ± 0.00 | 41.6 ± 1.07 | 90.6 ± 0.14 | 4.73 ± 0.19 | 5.47 ± 0.29 | 37.3 ± 2.01 | 19.3 ± 1.65 |
| 6 h | | 2,693 ± 122.6 | 8.50 ± 0.08 | 40.7 ± 2.24 | 91.2 ± 0.20 | 10.8 ± 2.06 | 4.90 ± 0.08 | 41.9 ± 2.11 | 12.9 ± 0.59 |
| 12 h | | 2,593 ± 66.0 | 8.57 ± 0.12 | 39.1 ± 0.85 | 90.5 ± 0.21 | 16.5 ± 3.14 | 5.60 ± 0.41 | 39.7 ± 0.76 | 9.0 ± 0.85 |
| 10 m | 9 | 2,700 ± 149.7 | 8.47 ± 0.05 | 38.4 ± 4.69 | 91.1 ± 0.06 | 4.30 ± 0.30 | 7.00 ± 0.22 | 39.2 ± 2.59 | 38.5 ± 4.71 |
| 3 h | | 2,693 ± 183.5 | 8.37 ± 0.17 | 41.8 ± 4.60 | 90.6 ± 0.17 | 8.77 ± 1.11 | 7.03 ± 0.19 | 43.0 ± 1.46 | 41.9 ± 2.71 |
| 6 h | | 2,827 ± 92.9 | 8.50 ± 0.08 | 41.5 ± 1.78 | 90.8 ± 0.68 | 13.7 ± 1.21 | 5.87 ± 0.17 | 37.9 ± 1.76 | 22.9 ± 1.35 |
| 12 h | | 2,753 ± 57.3 | 8.40 ± 0.08 | 38.9 ± 2.68 | 90.6 ± 0.20 | 37.1 ± 12.7 | 4.70 ± 0.54 | 37.7 ± 2.24 | 15.4 ± 1.52 |

Average ± standard deviation (n = 1–3). Conc.; Concentration. Con.; Experimental control. N.D.; Not detected. [*]Solvent control containing 0 or 9 mg/L LAS with no-fish at 10 min. [†]One of three samples in LAS 0 mg/L showing 7.7 and for others not detected.
